# Large-scale phenotypic and genomic characterization of *Listeria monocytogenes* susceptibility to quaternary ammonium compounds

**DOI:** 10.1101/2023.09.07.556668

**Authors:** Mirena Ivanova, Martin Laage Kragh, Judit Szarvas, Elif Seyda Tosun, Natacha Friis Holmud, Alexander Gmeiner, Corinne Amar, Claudia Guldimann, TuAnh N. Huynh, Renáta Karpíšková, Carmen Rota García, Diego Gomez, Eurydice Aboagye, Andrea Etter, Patrizia Centorame, Marina Torresi, Maria Elisabetta De Angelis, Francesco Pomilio, Anders Hauge Okholm, Yinghua Xiao, Sylvia Kleta, Stefanie Lueth, Ariane Pietzka, Jovana Kovacevic, Franco Pagotto, Kathrin Rychli, Irena Zdovc, Bojan Papić, Even Heir, Solveig Langsrud, Trond Møretrø, Roger Stephan, Phillip Brown, Sophia Kathariou, Taurai Tasara, Frank Aarestrup, Patrick Murigu Kamau Njage, Annette Fagerlund, Lisbeth Truelstrup Hansen, Pimlapas Leekitcharoenphon

## Abstract

*Listeria monocytogenes* is a significant concern for the food industry due to its ability to persist in the food processing environment. Decreased susceptibility to disinfectants is one of the factors that contribute to the persistence of *L. monocytogenes*. The objective of this study was to explore the diversity of *L. monocytogenes* susceptibility to quaternary ammonium compounds (QACs) using 1,671 *L. monocytogenes* isolates. This was used to determine the phenotype-genotype concordance and characterize genomes of the QAC sensitive and tolerant isolates for stress resistance, virulence and plasmid replicon genes. Distribution of QAC tolerance genes among 37,897 publicly available *L. monocytogenes* genomes were also examined. The minimum inhibitory concentration to QACs was determined by the broth microdilution method and non-sequenced isolates (n=1,244) were whole genome sequenced. Genotype-phenotype concordance was 99% for benzalkonium chloride, DDAC and a commercial QAC based sanitizer. Prevalence of QAC tolerance genes was 23% and 28% in our *L. monocytogenes* collection and in the global dataset, respectively. *qacH* was the most prevalent gene in our collection (61%), with 19% prevalence in the global dataset. Notably, *bcrABC* was most common (72%) globally, while 25% in our collection. Prevalence of *emrC* and *emrE* was comparable in both datasets, 7% and 2%, respectively. Replicon genes, indicative of plasmid harborage, were detected in 44% of the isolates and associated with the QAC tolerant phenotype. The presented analysis is based on the biggest *L. monocytogenes* collection in diversity and quantity for characterization of the *L. monocytogenes* QAC tolerance at both phenotypic and genomic levels.

**IMPORTANCE:** Contamination of *Listeria monocytogenes* within the food processing environment is of concern to the food industry due to challenges in eradicating the pathogen once it becomes persistent in the environment. Genetic markers associated with increased tolerance to disinfectants have been identified, which alongside factors favor the persistence of *L. monocytogenes* in the production environment. By employing a comprehensive large-scale phenotypic testing and genomic analysis our study significantly enhances the understanding of the prevalence of quaternary ammonium compound (QAC) tolerant *L. monocytogenes* and the genetic determinants associated with the increased tolerance. Furthermore, we report on the prevalence of QAC tolerance genes among 37,897 publicly available *L. monocytogenes* sequences and their distribution within clonal complexes, isolation sources and geographical locations. As the propagation of QAC tolerance showed not be evenly distributed globally this highlights that understanding the development of *L. monocytogenes* disinfectant tolerance can be monitored using publicly available WGS data.

## INTRODUCTION

*Listeria monocytogenes* is a free-living saprophyte and an intracellular pathogen responsible for listeriosis, a serious foodborne infection with life-threatening complications in “at-risk populations” such as individuals with weakened immune system, elderly, neonates, as well as miscarriages in pregnant women (1). Unlike campylobacteriosis and salmonellosis, which are the two leading bacterial causes of foodborne illnesses, listeriosis has a low incidence rate but a high mortality rate (2). Contaminated ready-to-eat (RTE) foods have been identified as the major cause of several large outbreaks, including foods such as polony in South Africa (3), cold cuts in Canada and Denmark in 2008 and 2014, respectively (4,5), and cheese made with unpasteurized milk in USA (6).

Food contamination may occur in the food production facilities where *L. monocytogenes* can enter with e.g., raw materials, personnel or equipment and get established and persist for decades in the food processing environment (FPE) despite ongoing cleaning and disinfection programs (7), or it can be constantly introduced into FPEs with incoming raw materials. Both biotic (e.g., biofilm formation ability, growth at refrigeration temperatures, increased tolerance to sanitizers and disinfectants, heavy metals and desiccation, etc.) and abiotic (e.g., improper cleaning, biocide residuals, poor hygienic design, difficult to clean areas, etc.) factors are involved in successful survival and persistence of *L. monocytogenes* in FPEs, likely leading to contamination with *L. monocytogenes* clones with increased persistence potential (8). Some *L. monocytogenes* clonal complexes (CCs) have been established as successful persisters within FPEs, such as CC121 and CC9 that are also associated with hypovirulent strains, while others have been strongly implicated with clinical sources and hypervirulence (CC1, CC2, CC4 and CC6) (9)). Certain hypervirulent CCs (e.g., CC2) can also persist in FPE (Guidi et al., 2021).

To help control *L. monocytogenes* in FPEs, effective and robust cleaning and disinfection programs are needed. Within cleaning and disinfection programs, three major biocide (disinfectants or sanitizers according to EU and US definition, respectively) categories are widely applied in the food industry, namely quaternary ammonium compounds (QACs) with main representatives benzalkonium chloride (BC) and didecyldimethylammonium chloride (DDAC); peroxygens, (e.g., peracetic acid (PAA) and hydrogen peroxide); and halogen-releasing agents, such as sodium hypochlorite (NaClO) (13–15). To date, few specific genetic determinants associated with increased tolerance to oxidative agents such as PAA and NaClO have been described, possibly due to the multiple modes of action of these agents (14,16,17). In contrast, genetic determinants associated with increased tolerance to QACs have been reported and include the efflux pumps genes *bcrABC, emrC*, *emrE* and *qacH* that provide *L. monocytogenes* with increased survival in the presence of low concentrations of biocides (18–21). The minimum inhibitory concentrations (MICs) of the QAC-based disinfectants towards *L. monocytogenes* have been reported to vary between studies, ranging from 0.6 to >20 mg/L (17,22–25), while the recommended concentrations in FPEs are 200–1000 mg/L facilities (27). Therefore, it has been accepted that the term “tolerance” is more appropriate to describe the reduced *L. monocytogenes* susceptibility to disinfectants than “resistance” (26).

Reduced *L. monocytogenes* susceptibility to food industry disinfectants, specifically to QACs, has been the focus of many studies (17,22,27–29), including co-occurrence with antibiotic resistance genes (22,30). However, the lack of a standardized disinfectant susceptibility method does not allow comparison between the results reported by these studies. Variations in methods and factors such as inoculum level, disinfectant dilutions and concentrations, media used, and incubation conditions can contribute to variability of the MIC values without having a real connection to genetic variation. At the same time, various efflux pump genes have been shown to play a role in increased tolerance to QAC disinfectants in *L. monocytogenes*, including the *bcrABC* cassette, *emrE*, *emrC*, *qacH* (carried by Tn*6188*), *mdrL*, and *lde* genes (31–33). Here, we aimed at elucidating the genotype-phenotype concordance by performing disinfectant susceptibility testing of 1,671 *L. monocytogenes* isolates to two QAC substances, BC and DDAC, and a commercial QAC based disinfectant and to whole-genome sequence 1,244 isolates that have not been previously sequenced. We analyzed the genomes of the QAC sensitive and tolerant isolates for stress resistance genes, virulence factors and plasmid content. Additionally, the global dissemination of *bcrABC*, *qacH*, *emrC* and *emrE* genes, which were found exclusively in the tolerant isolates in this study, was examined in 37,897 publicly available raw sequencing data, deposited in the European Nucleotide Archive (ENA) as of April 2021. The distribution of QAC-associated genes in the global dataset among CCs, isolation sources and geographical locations was also investigated.

## RESULTS

### Characterization of the *L. monocytogenes* strain collection and QAC tolerance

To capture the full potential diversity of *L. monocytogenes* to QACs isolates from twenty collaborating laboratories were combined to create a diverse collection of isolates in relation to isolation year, environmental and geographical origin. In total 1,671 *L. monocytogenes* were included from >19 countries collected within a time span of 98 years, from 1924 to 2021 with >100 isolates from before 1990 (Fig. 1, Table S1). The isolates were of different origin with isolates sampled from human (n = 83; 5%), animal (n = 122; 7%), food (n = 839; 50%), FPEs (n = 488; 29%), feed (n = 4; <1%), natural environment (n = 32; 2%), farm environment (n = 66; 4%) and unknown sources (n = 22; 1%) (Fig. 1A). Isolates from different environments and countries were included as the stresses (environmental, antimicrobial, human etc.) faced by *L. monocytogenes* vary, which could affect the QAC tolerance and adaptation of the isolates from different niches and countries (Fig 1B). All *L. monocytogenes* isolates from collaborating partners were included in the strain collection and screened for QAC tolerance before whole genome sequencing. Minimum inhibitory concentration (MIC) towards BC were determined for all isolates and 1283 isolates (77%) were defined as BC sensitive isolates with a MIC < 1.25 mg/L, while 388 isolates (23%) were defined as BC tolerant with MIC > 1.25 mg/L.

**Figure 1.**
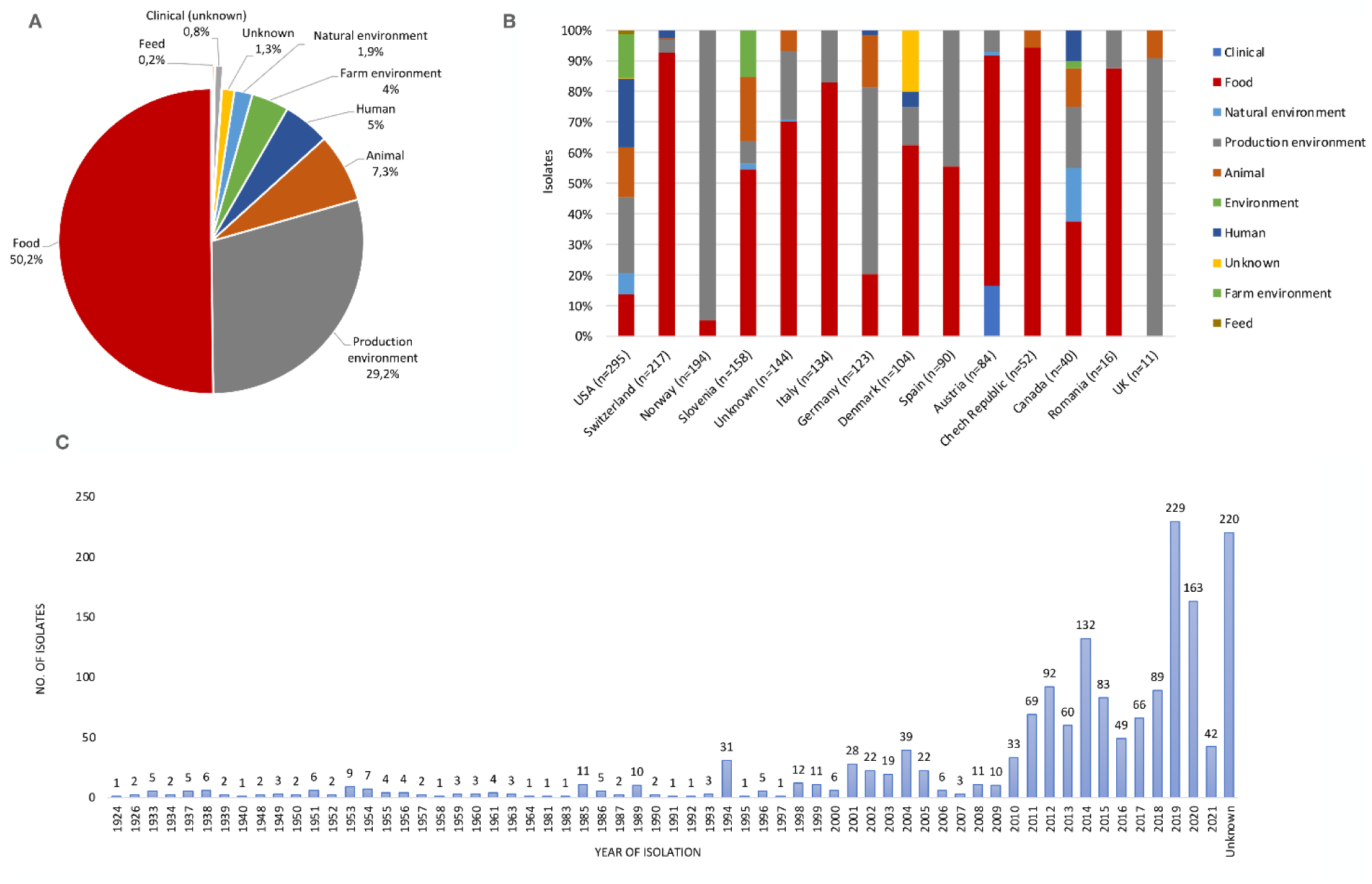
**A.** Percentage of *L. monocytogenes* isolates by source of isolation. Sources below 0.1% are not shown. **B.** Division of isolates by source of isolation for each country of collection. Countries with < 10 isolates are not shown on the figure: Turkey (n=3, food), Russia (n=3, food), Belgium (n=1, food), China (n=1, food), Finland (n=1, food) and France (n=1, production environment). **C.** Temporal distribution of the isolates in the collection covering a time span of 98 years.

### Genomic characteristics and phylogenetic analysis of the *L. monocytogenes* isolates

Genomes of the 1,671 *L. monocytogenes* isolates ranged from 2.75 to 3.32 Mbp in size with a G+C content of 37.7 to 38.3% (Table S2). The isolates were assigned to four phylogenetic lineages based on the genome alignment of 2,146 core genes present in ≥ 99% of the isolates – lineage I (LI, n = 589, 35%), lineage II (LII, n = 1064, 64%), lineage III (LIII, n = 17, < 1%) and lineage IV (LIV, n = 1, <1%) (Fig. S1). Within the isolate set, there were six serogroups (1/2a (n = 875, 52%), 1/2b (n = 262, 16%), 1/2c (n = 165, 10%), 4b (n = 319, 19%), IVb-v1 (n = 10, 0.6%), L (n = 19, 1%), atypical (n = 1, <1%) and undetermined (n = 20, 1.2%)), and 75 CCs and 20 singleton sequence types (STs). Eleven CCs predominated, accounting for 67% (n = 1,117) of all the examined isolates. The distribution of the sources of isolates for the 11 most predominant CCs is presented in Fig. 2.

**Figure 2.**
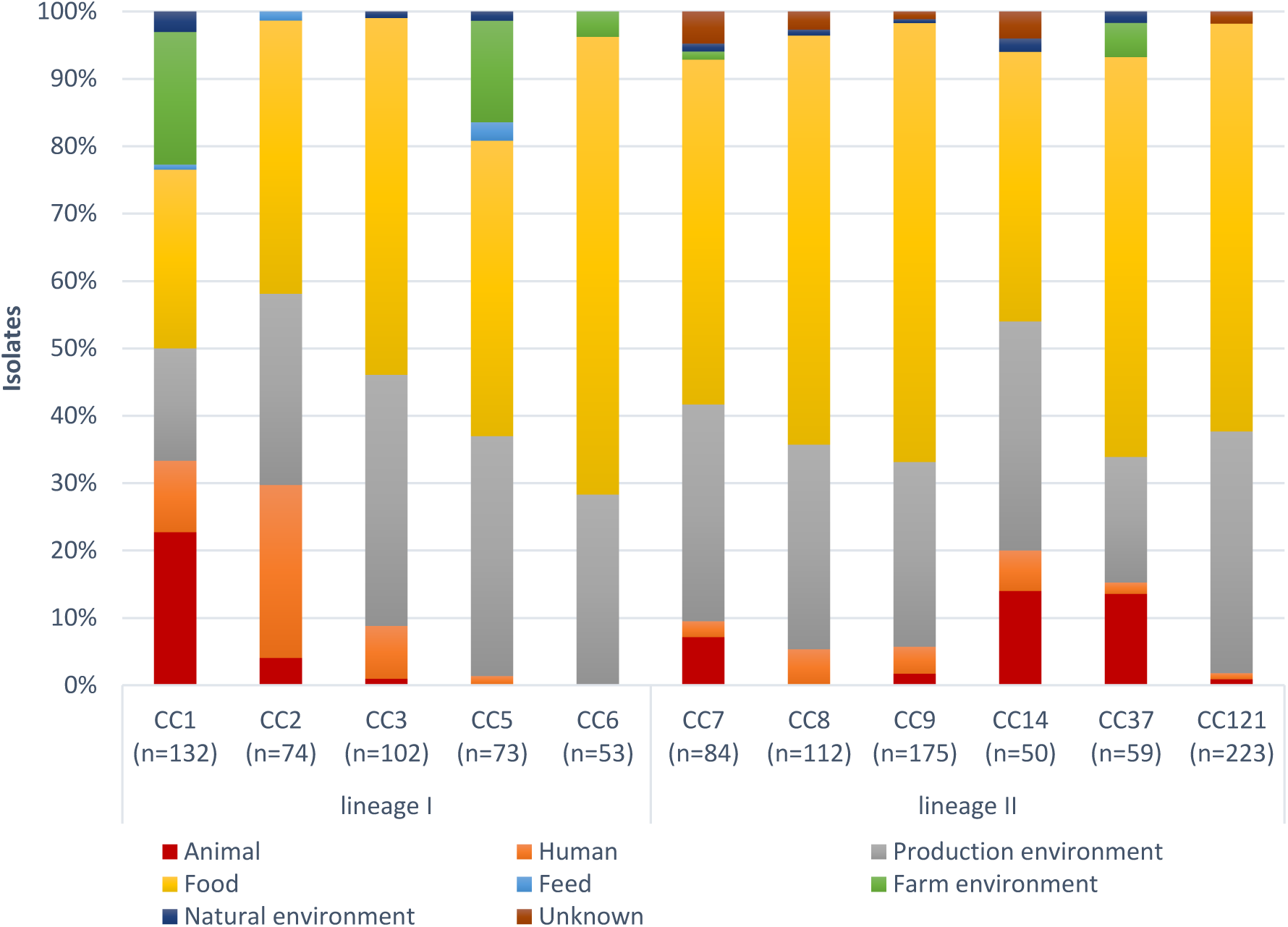
Distribution of the main isolation sources within dominating CCs (CCs with > 50 isolates are shown in the figure).

### Phenotype-genotype concordance of the *L. monocytogenes* QAC susceptibility

The genomes of the 1,671 *L. monocytogenes* isolates were screened for the presence of QAC tolerance genes (i.e., *bcrABC*, *emrC*, *emrE* and *qacH*) and the genotypes were associated with the respective phenotype (BC MIC values). The genes *bcrABC*, *emrC*, *emrE* and *qacH* were detected only in isolates with MIC ≥ 1.25 mg/L. Based on that, an MIC cut-off of ≥ 1.25 mg/L for BC tolerance was established (Fig. 3) and isolates were classified as BC sensitive (77%, n=1283) having MIC < 1.25 mg/L and BC tolerant (23%, n=388) with MIC ≥ 1.25 mg/L. Twenty (5%) of the BC tolerant isolates did not carry any of the four QAC tolerance genes associated with the tolerant phenotype. Moreover, no BC sensitive isolates with a QAC tolerance gene, nor BC tolerant isolates carrying more than one QAC tolerance gene were found. A randomly selected subset of isolates was tested for sensitivity to DDAC (n=247) and Mida San 360 OM (cQAC) (n=155), respectively. There was a clear distribution of the MIC values for DDAC and cQAC and the cut-offs for tolerance were drawn at MIC > 0.4 mg/L and ≥ 0.63 mg/L for DDAC and cQAC, respectively, based on presence/absence of the four QAC tolerance genes (Fig. 3). All BC tolerant isolates were also tolerant to DDAC and cQAC, except one isolate (414a-1), which was tolerant to BC but not to cQAC.

**Figure 3.**
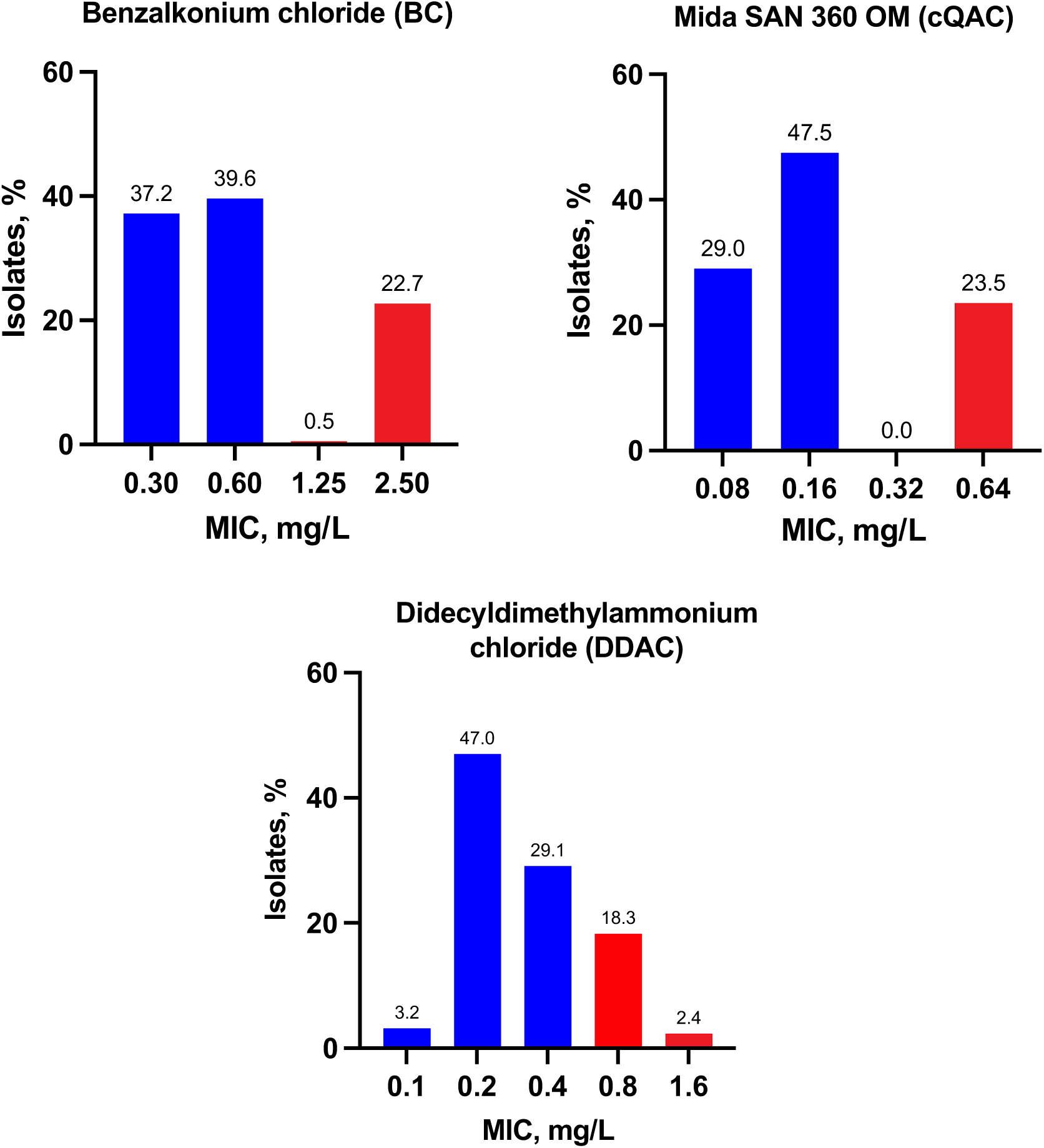
Distribution of the MIC values of sensitive (blue) and tolerant (red) *L. monocytogenes* isolates to Benzalkonium chloride (n=1671), Mida SAN 360 OM (cQAC, n=155) and Didecyldimethylammonium chloride (DDAC, n=247).

The most prevalent QAC tolerance gene found within our isolate collection was *qacH* (61% of the BC tolerant isolates; n=237), followed by *bcrABC* (25%, n=96), *emrC* (7%, n=27) and *emrE* (2%, n=8) (Fig. S2A). The majority of the *qacH*-harbouring isolates belonged to CC121 (83%) and CC9 (12%). The isolates carrying *bcrABC* genes most commonly belonged to CC9 (30%), CC5 (26%) and CC321 (16%). The *emrC*-harbouring isolates belonged to 11 CCs, and among them CC6 (41%), CC14 (11%) and CC403 (11%) were the most prevalent CCs. Meanwhile, *emrE* was detected among CC8 isolates. Notably, the BC-tolerant isolates without an identified QAC tolerance gene were genetically diverse, belonging to 12 different CCs (Fig. 4A). Overall, the majority of isolates harboring QAC tolerance genes belonged to LII (85%), with 60% and 20% belonging to CC121 and CC9, respectively. QAC tolerance genes were not detected among the LIII and LIV isolates. The majority of the QAC tolerance genes were found in isolates from food (56%) and FPE (38%), and only 2.6% were from clinical sources (Fig. 4B). All *bcrABC* and *emrC* genes were detected on plasmid contigs. The majority of the *qacH* genes (n = 237) were located on contigs with no identified replicon gene and with lengths greater than the largest identified *Listeria spp*. plasmid (>152 kbp, CP022021.1), assuming chromosomal origin.

**Figure 4.**
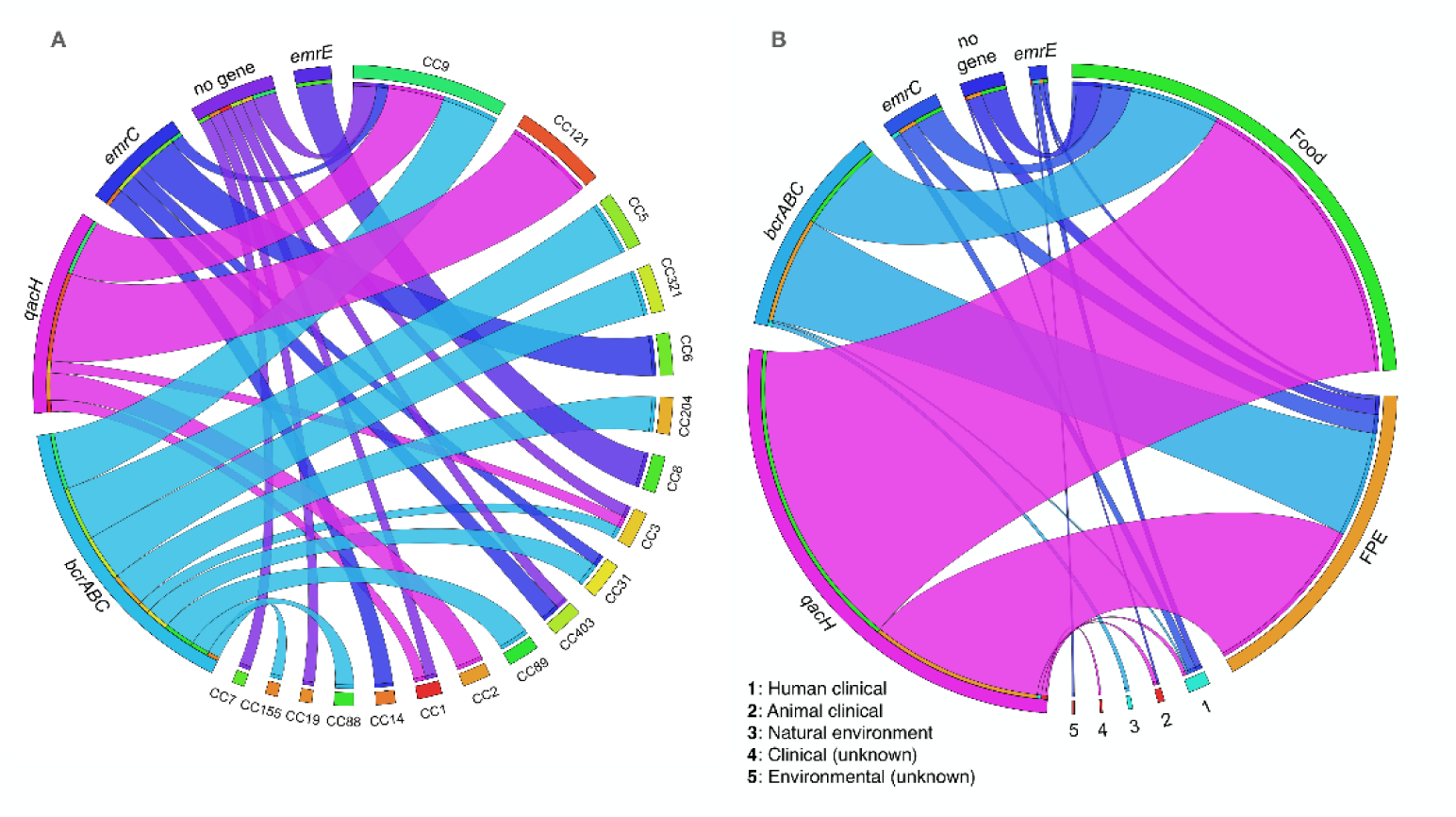
Genomic characterisation of the *L. monocytogenes* tolerance to QACs. Distribution of CCs **(A)** and isolation sources **(B)** among the isolates harbouring QAC tolerance genes and those isolates with unknown mechanism of tolerance. Graphs were generated by Circos Viewer v0.63-10 (http://mkweb.bcgsc.ca/tableviewer/).

### Identification of plasmids among the QAC tolerant and sensitive isolates

Genetic determinants of the QAC tolerance are often plasmid-borne (e.g., *bcrABC*, *emrC*), therefore an analysis to identify plasmid content in the tolerant and sensitive isolates was performed. Overall, 44% of the isolates (n=735) were determined to carry plasmids. Of these, 712 carried one replicon gene, 24 carried two replicon genes, and three carried plasmids with unknown replication system. Ninety-seven percent of the plasmid-harboring isolates (714/735) carried *repA* genes. In total 728 *repA* genes were subjected to a phylogenetic analysis, presented in Fig 5A. The majority of the *repA* genes belonged to phylogenetic groups G1 (n=326; 20%) and G2 (n=390; 23%), and 10 genes were allocated to G4. Two *repA* variants clustered independently of any known RepA group and were most closely related to G3; they were named G13 (Fig. 5A). These two G13 isolates (SKB398 and SKB102) belonged to CC912 and ST1365, originating from black bears. Additionally, 20 and three of the 735 isolates with identified plasmids carried *repB* genes and plasmids with an unknown replication system, respectively. The *repB* genes were found on 4 kb contigs, except for isolates N195 and LIS08. In these two isolates, the *repB* genes were co-located with *repA* genes on plasmid contigs that were of length 53 and 81 kbp, respectively. The majority of the *repA* genes were identified in isolates from food (55%) and FPE (39%), while only 3.4% and 0.8% were seen in clinical, and farm and natural environment isolates, respectively (Fig. 5A). Food and FPE isolates were evenly distributed among the plasmid phylogenetic groups in this study. Prevalence of isolates determined as sensitive or tolerant to QAC were in addition not significantly more associated with food or FPE as only significant differences were seen for human clinical samples where QAC sensitive isolates were more prevalent (Fig. 5B). It is worth mentioning that 91% of the *repA* genes in the phylogenetic group G2 belonged to LII isolates, while G1 was equally distributed within LI and LII. Notably, 98% of the CC121 isolates carried *repA* genes belonging to G2, while G1 was the predominant phylogenetic group (88%) in the CC9 isolates. Similarly, other CCs with high frequency of *repA* had different distributions of the phylogenetic groups with CC3 (91% G1), CC5 (50% G1, 50% G2) and CC8 (96% G2. *repB* genes showed the highest nucleotide identity and coverage to pLmN12-0935 (CP038643.1). Two of the three plasmid contigs with unknown replication systems were most similar to plasmid pLIS55 (MZ151539.1), while the *repB* gene in isolate CDL77 had no match to any known sequence in NCBI. The majority of the isolates which carried simultaneously two *repA* genes came from FPEs and belonged to CC3, CC5, CC8 and CC89.

**Figure 5.**
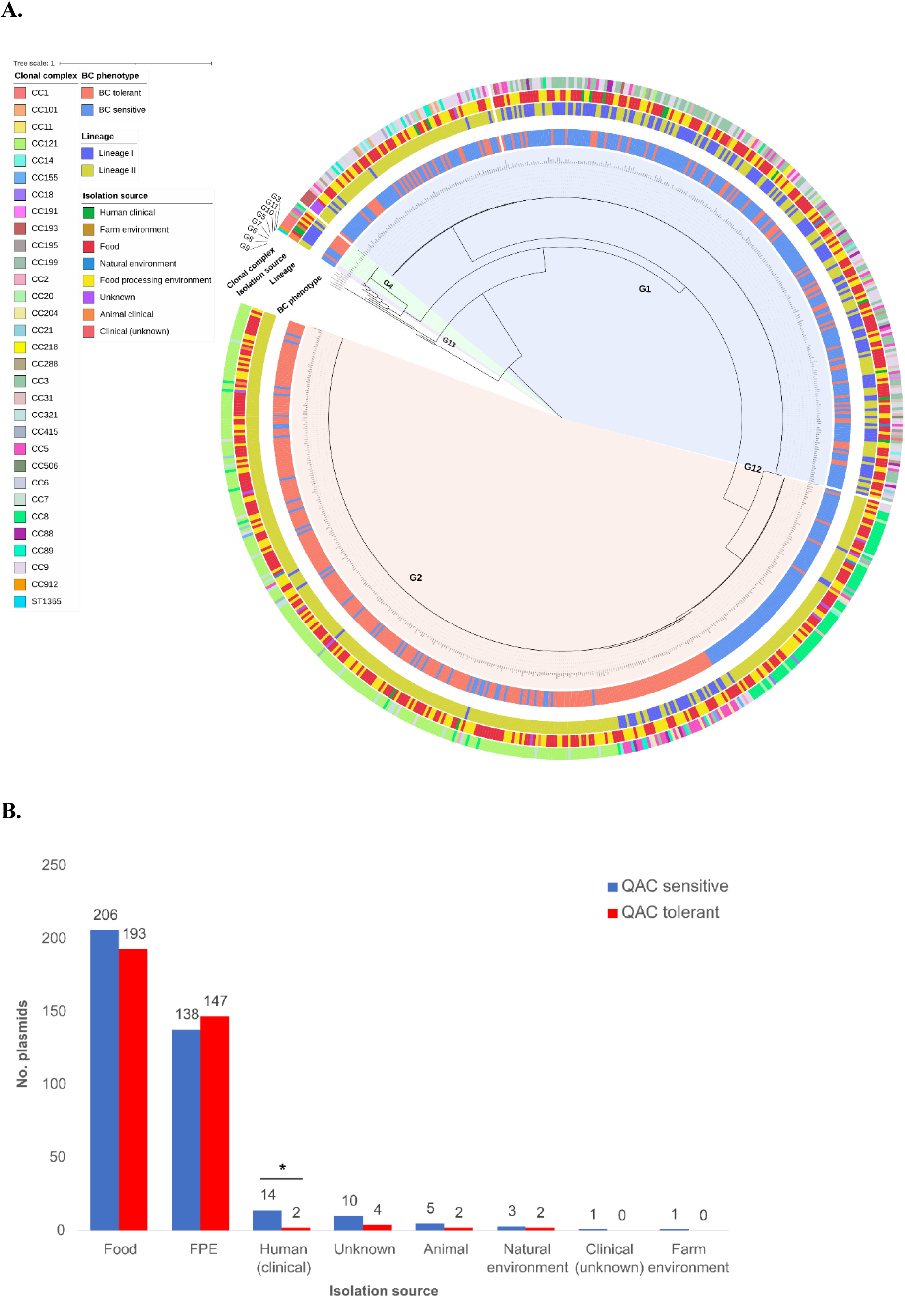
**A.** Mid-point rooted ML phylogenetic tree of the RepA sequences identified in 728 of the *L. monocytogenes* isolates. The 13 plasmid replicon groups detected among the isolates in this study and the reference groups are indicated. **B.** Prevalence of plasmid carrying isolates determined as sensitive or tolerant to QAC and their isolation source. Asterisks indicate significant differences determined with the Pearson’s chi-squared association test with p < 0.05.

Further, blastn screening using a custom database of *L. monocytogenes* virulence and stress resistance genes against the contigs carrying replicon genes (n = 735) identified co-location of *qacH* and *repA* genes in six of the G4 plasmid group isolates. In these isolates, *qacH* had a different genetic context from the previously described Tn*6188* transposon structure, consisting of *tetR* and *qacH* genes, and an upstream transcriptional regulator *mutR* (Fig. S3).

All *bcrABC* and *emrC* genes were detected on plasmid contigs as determined by their co-location with other plasmid-associated genes. While the *bcrABC* gene was located on contigs of varying sizes (3.8 – 91.2 kbps), the *emrC* was found on small contigs (4.4 - 4.7 kbps) with higher nucleotide coverage compared to the coverage of the other contigs in the assemblies. The *qacH* genes (n=237) were located on contigs with no identified replicon gene and with lengths greater than the largest identified *Listeria* spp. plasmid (>152 kbp, CP022021.1), suggesting chromosomal origin. *emrE* as part of the LGI-1 was located on the chromosome in all isolates as previously described (18).

The LIPI-1 genes, *plcA*, *hly*, *mpl*, *plcB* and a truncated *actA* gene, associated with increased *L. monocytogenes* virulence, were also found co-located with a *repA* gene in two isolates (ERR1432982 and ERR1432994). Genes encoding for resistance to arsenic (*ACR3*, *lmo2230*, *arsA*, *arsD*, *arsR*) (n = 8), cadmium resistance genes *cadA1C1* (n= 522) and *cadA2C2* (n = 45), heat resistance gene *clpL* (n = 348), genes for copper resistance *copB* (n = 198) and *copL* (n = 54), mobile genetic element consisting of *gbuC* (n = 173), NiCo riboswitch (n = 168) and *npr* (n = 173), *tmr* (n=20), *mco* (n = 197) and the mercury resistance operon (n = 2) were also co-carried with a *repA* gene (Table S6).

### Genetic organisation of the *bcrABC*-harbouring plasmid contigs

The *bcrABC* gene was detected in 96 isolates representing 11 CCs and four isolation sources (Fig. S4). Using multiple sequence alignment, the *bcrABC*-carrying contigs were grouped into eight structures (Fig. S5). Structures 1 and 1a, detected only in CC9 isolates, were identical except for the presence of the genetic module NiCo riboswitch-*gbuC*-*npr* that was seen in structure 1 but absent from 1a. Structures 2, 3 and 4 had a common backbone consisting of *ltrC*, *bcrABC*, recombinase, glyoxalase, *tmr*, *qorB* and *ravA*. While structure 2 was detected only in the CC5 231a1 isolate, structures 3 and 4 were distributed among various CCs. A shared feature of the structures 5, 5a and 6, found in three non-clonal *L. monocytogenes* isolates from CC1 (LI),) and CC31 and CC204 (LII), was the presence of the mercury resistance (*mer*) operon. Blastn search in NCBI of the *bcrABC* contigs of structures 5 and 5a identified similarity to pLMR479a (HG813248.1) from a CC8 isolate from smoked salmon, with 84% coverage, lacking only the *bcrABC* cassette and the *mer* operon, suggesting that these genetic elements could horizontally be transferred together. The isolate with structure 6, N21-0102 isolated from poultry in Switzerland, had 99.89-100% identity and over 100% coverage to the *Listeria innocua* plasmids CP095722.1 and CP095729.1, found in isolates from meat in Jamaica. The sequences from the three CC31 isolates from milk in Switzerland with structure 7 had 100% coverage and 100% identity to a *L*.

*innocua* pLIS35 plasmid (MZ127844.1), isolated from a food contact surface swab in Poland. Structure 8 was detected only in CC321 isolates from North America, in which the *bcrABC* cassette was carried on a composite transposon associated with IS*1216* (Fig. S5). This shows that there is a wide diversity of genetic contexts surrounding the *bcrABC* operon, although it seems that the cassette is always located on a plasmid.

### Identification of virulence and stress resistance genes among QAC tolerant and sensitive isolates

The carriage of virulence factors and stress resistance genes have previously been linked to specific CCs, environments and QAC phenotypes in small- and large-scale screening studies. To validate and extent these observations with a larger isolate collection a variety of virulence and stress resistance genes were identified in the 1,671 *L. monocytogenes* isolates.

Ten different mutations (including four newly detected in this study) and two internal deletions in the *inlA* gene, leading to premature stop codons (PMSCs) and proteins with various lengths, were found in 21% of the isolates (Table S5). The majority of these isolates belonged to lineage II (97%) and specifically CC121 (61%) and CC9 (30%). The most common mutation (C -> T at nucleotide 1474 leading to a protein with 491 aa), accounting for 61% of the total mutations, was only seen in CC121 isolates (34). All but five CC121 isolates (98%) carried an *inlA* PMSC mutation. Truncated *inlB* genes, with deletion of amino acids from 356/357 to 630 (leading to a protein of 356/357 aa), were detected in 19 CC5 isolates originating from FPEs and food samples. PMSCs in *inlB* gene resulting in shorter protein sequences were identified in five food isolates belonging to CC1 and CC8 (data not shown). All isolates with truncated *inlB* genes harboured intact *inlA* genes. The *actA* gene, located on LIPI-1, was truncated in all isolates from 20 CCs, including CC1, CC121, CC31, CC4, CC379 and CC88 (Fig. 6A).

**Figure 6.**
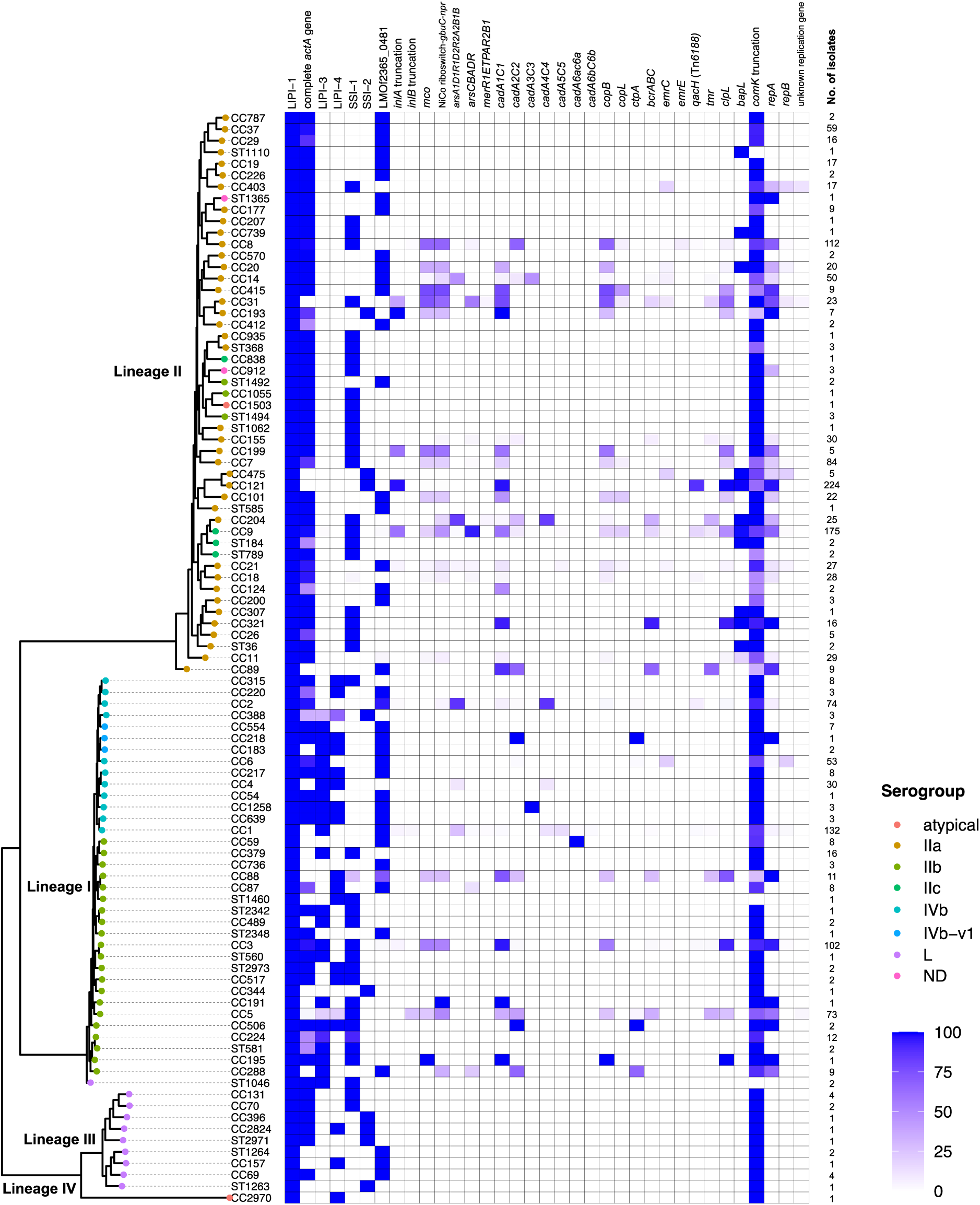
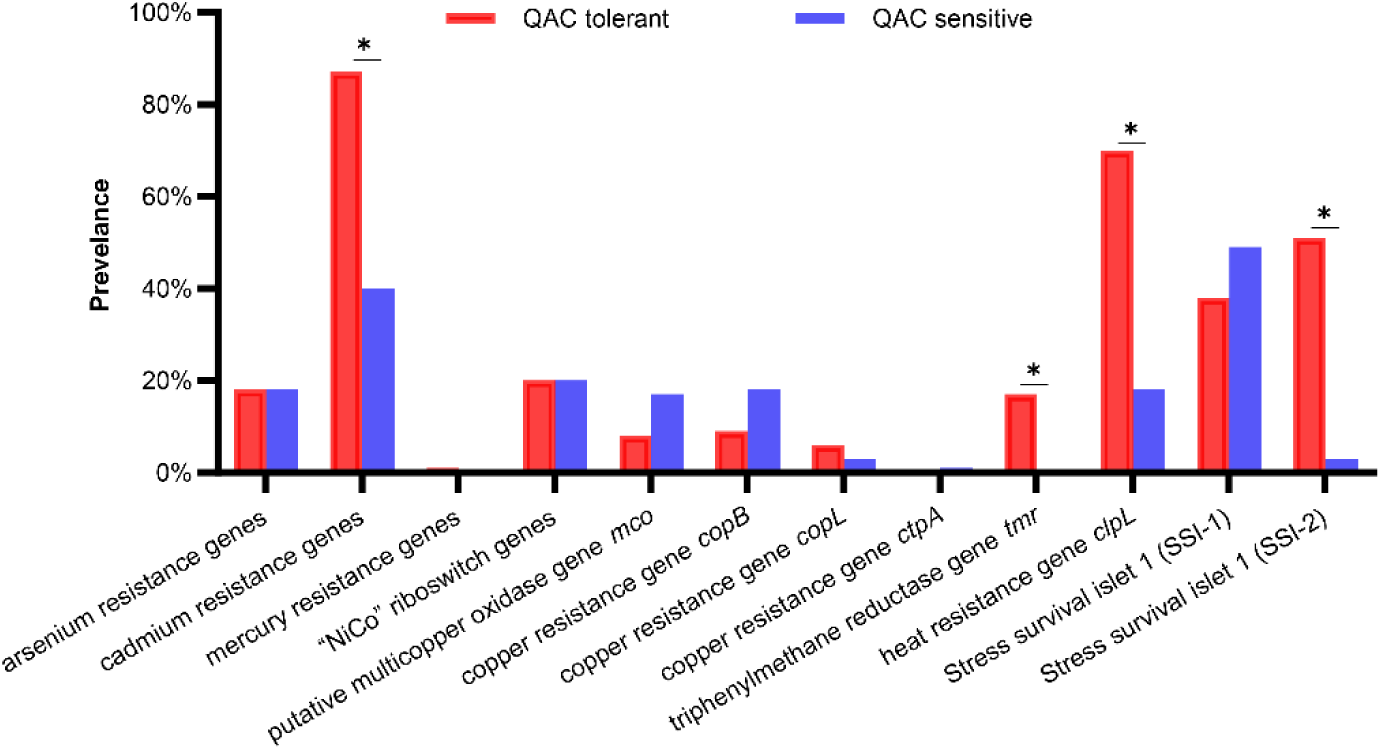
**A.** Stress resistance, virulence and plasmid replicon gene prevalence among the 95 CCs and singleton STs in this study presented as percentage within each CC. A gene is considered present if the gene coverage is > 90%. The phylogeny is a core-genome ML tree built with a representative isolate (the most recent ancestor) from each CC with serogrouping based on the serogrouping scheme from https://bigsdb.pasteur.fr/_nuxt/img/serogroups.9daaa98.png **B.** Prevalence of stress resistance genes among the QAC tolerant (n=388) and sensitive (n=1283) isolates. Asterisks indicate significant differences in prevalence determined with the Pearson’s chi-squared association test with *P* < 0.05.

The cadmium resistance genes *cadA1C1* and *cadA2C2*, located on plasmids with various sizes, were the predominant cadmium resistance genes, while *cadA3C3*, *cadA4C4*, *cadA5C5*, *cadA6aC6a*, *cadA6bC6b* were less frequently detected. Copper (*mco*, *copB*, *copY*, *zosA*, *ctpA*), arsenic (*arsA1D1R1D2R2A2B1B2* and *arsCBADR*) and mercury (*merR1ETPAR2B1*) resistance genes were unevenly distributed (Fig. 6A). The oxidative stress, heavy metal, salt and acid resistance cassette NiCo riboswitch-*npr*-*gbuC*-like, were transferred together by the same mobile genetic element and found in CCs that carry plasmid genes. The environmental stress islet SSI-1 was present in 45% of the CCs, while SSI-2 was less prevalent (9%). The gene *comK*, a common hotspot for prophages in *L. monocytogenes*, was truncated in 82% of the isolates, and a truncation was found in all but three CCs. Two insertion sites were identified, TAA-TAAAA (at nucleotides 212 and 214) seen among 25% of the isolates, and GGA (at nucleotide 190) seen in 57% of the isolates. Additionally, 98.6% of the isolates contained full or partial dsDNA phage sequences and 52.3% had full phage sequences as identified by VirSorter with minimum length of 1500 bp. The *bapL* gene, associated with biofilm formation, was present in CCs from LII only. The LIPI-1 pathogenicity island, that carries the master regulator for virulence transcription (*prfA*), and cluster of virulence genes used to escape from vacuolar compartments (*hly*, *plcA*; *actA*; *mpl*, *plcB*, and *orfX*), was present in all isolates. The pathogenicity islands LIPI-3 and LIPI-4 were present in the majority of the isolates in CCs from LI and were absent in isolates from LII. CC218, CC217, CC506, CC1258 and CC639 harboured all LIPI-1, -3, and -4 genes, indicative of potential to exhibit higher virulence level. The isolates from LIII and LIV did not carry any of the screened stress resistance genes (Fig. 6A, Table S7). When isolated were grouped as sensitive and QAC tolerant there were differences in the prevalence of several stress resistance genes between the groups. When adjusting for the higher number of sensitive isolates in the isolate collection there were a significantly (*P*<0.05) higher prevalence in QAC tolerant isolates of genes conferring resistance to cadmium, and heat as well as SSI-2 (Fig. 6B).

### Global distribution of QAC tolerance genes in *L. monocytogenes*

The raw sequencing data from 39,196 *L. monocytogenes* isolates deposited in the European Nucleotide Archive (ENA) were screened for the presence of QAC-tolerance genes *bcrABC*, *emrE*, *emrC* and *qacH* (Table S8). Of the analyzed sequencing data, 10,953 (28%) carried one or more QAC tolerance genes. *bcrABC* was the most abundant gene, present in 72% of the QAC tolerance gene positive isolates, followed by *qacH* (19%), *emrC* (7%) and *emrE* (2%) (Fig. S2B). *qacH* had the highest nucleotide variability among the four QAC tolerance genes, and nine *qacH* variants with nucleotide identity above 90% were detected in the global dataset based on analysis of raw reads using KMA (Fig. S6). Notably, 23 isolates carried simultaneously two QAC tolerance genes. Since several large-scale studies using in-house sequencing data (35,36), including this study, detected no more than one QAC tolerance gene in a single *L. monocytogenes* genome, the raw sequencing data of the isolates carrying two QAC tolerance genes were assembled, re-screened by blastn and sub-typed by MLST. Subsequently, 7/23 isolates carried only one QAC gene. Of the 16 isolates that still carried two QAC genes, two alleles of an MLST gene were detected in three isolates, suggesting contamination of the sequencing reads. For the remaining 13 genomes, in which *bcrABC* and *qacH* (seven isolates), *bcrABC* and *emrC* (one isolate), and *qacH* and *emrC* (four isolates) were simultaneously detected, re-isolation and re-sequencing could reveal if two QAC tolerance genes could in fact be carried by the same *L. monocytogenes* isolate (Table S9). All QAC tolerance genes were present in *L. monocytogenes* isolates from LI and LII, with the exception of one LIII (ST1142) isolate (SRR8223359) that harbored *bcrABC* and recovered from a FPE isolate in the United States.

There were differences in the dissemination of the QAC tolerance genes among countries/continents and CCs. The *bcrABC* cassette was widely distributed in the United States, while *qacH* and *emrC* were mainly associated with Europe, and *emrE* had highest occurrence in Australia/Oceania (Fig. 7, Fig. S7, Fig. S8). The *emrC*, *emrE* and *qacH* were associated with CC8, CC6 and CC121, respectively, while the *bcrABC* cassette was distributed among several CCs, having the highest occurrence in CC5 followed by CC321, CC155, CC9 and CC7 (Fig. S9). All genes were associated with FPE or food and feed sources, except *emrC*, which was also associated with clinical sources (Fig. S10,).

**Figure 7.**
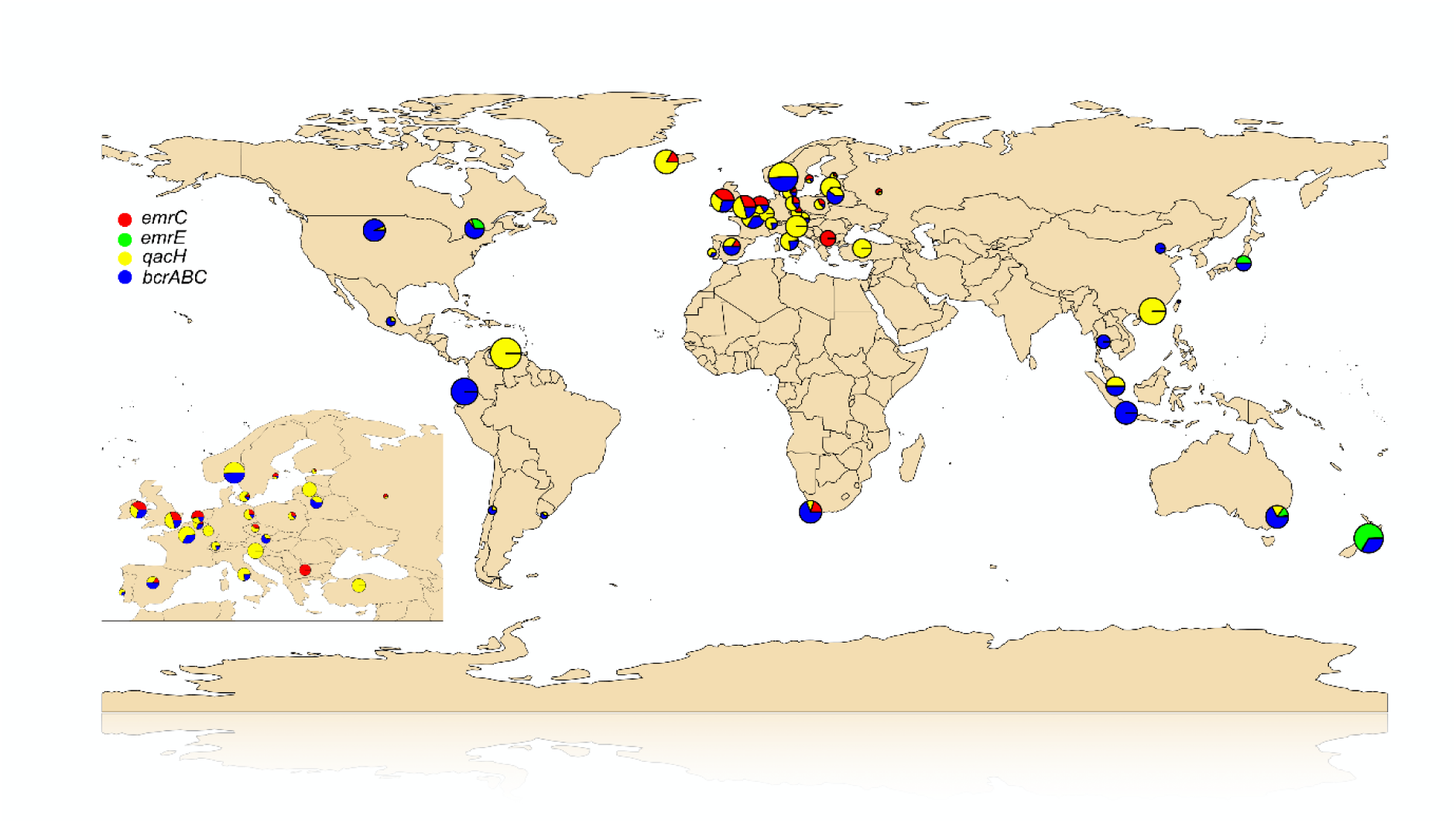
Distribution of *bcrABC*, *qacH*, *emrE*, *emrC* genes globally **(A)** and in Europe **(B)** in *L. monocytogenes* sequencing runs deposited in ENA as of 21 April 2021. The pie charts indicate the proportion (rate) of the genes in each of the country in which at least one of the genes was present and their size reflects the rate of each gene per 1000 genomes (overview in Table S11).

## DISCUSSION

The adaptation and persistence of *L. monocytogenes* in FPEs is a complex issue, often dependent on a combination of factors, such as biofilm formation capacity of different strains, improper equipment design, the presence of organic matter, insufficient removal of disinfectants during cleaning and disinfection and presence of genes conferring tolerance to various environmental stressors, among others (8,37). The QAC genes conferring tolerance to low concentrations of QACs can in combination with the above events contribute to persistence of *L. monocytogenes* as isolates harboring such genes have potentially higher survivability in the presence of low disinfectant concentrations in niches and often carry additional stress genes as seen in present study (Fig. 6B).

*L. monocytogenes* susceptibility to food industry disinfectants, specifically QACs, which are applied in the food industry, has been extensively examined (17,22,27,28,31,38). Most studies have included either a small number of isolates or isolates restricted to certain geographical areas, or focused on isolates recovered from specific environments, e.g., meat (24,39) or pork processing environments (40). The 1,671 *L. monocytogenes* isolates in this study were collected to represent diverse origin sources and geographical locations. Of all the isolates, 74% were selected solely based on the available metadata, therefore minimizing the bias of choosing isolates with certain genetic characteristics. Yet, *L. monocytogenes* isolates originating from FPEs and food prevailed in our collection, which could be based on sampling bias, as FPEs and foods make up the important niches for studying *L. monocytogenes* transmission routes (35,36,41,42).

Unlike for antibiotics, the lack of a harmonized assay for disinfectant susceptibility testing affects the comparability of the phenotypic results produced by different studies. The previously reported QAC MIC values vary significantly between studies due to differences in the biocide and/or inoculum concentrations, variations in media and assay conditions (27,28,29,54,55). For instance, *L. monocytogenes* exhibits different physiological traits at lower temperatures (e.g., 15°C) as compared to 37°C, such as the different types of motility and having different membrane fatty acid composition (43). An important aspect of our large-scale study is the consistency and reliability of the phenotypic data obtained using the same antimicrobial susceptibility method and conditions, which allowed us to establish a cut-off for tolerance based on the presence/absence of known genetic determinants. The observed distribution of the QAC MIC values allowed for a clear division of the sensitive and tolerant phenotypes. The high concordance between a tolerant phenotype and the presence of a QAC tolerance gene (99%) means that the phenotype of QAC tolerance can be predicted *in-silico* if the genotype is known without the need for conducting wet lab phenotypic testing.

The few tolerant *L. monocytogenes* isolates with no known QAC genetic determinants belonged to various CCs. This suggests a spread of mobilizable genetic elements distributed among diverse CCs, or that these isolates may be tolerant due to altered expression regulation of efflux pump genes, such as *mdrL* or *lde*. Other recent studies also reported QAC tolerant *L. monocytogenes* isolates with no known QAC tolerance genes, e.g., a food environment ST155 isolate from the United States (44) and two isolates from German food production facilities (22). Genome-wide association studies (GWAS) have the power to identify novel genetic determinants associated with disinfectant tolerant phenotypes using large collections of isolates (17). Among the pangenome orthologous groups (POGs), in the study of Palma et al. (17), Tn*6188*_*qacH* and a cluster of 14 POGs, a part of a prophage region, were strongly associated with BC tolerance (17). However, such findings have not yet been confirmed phenotypically. It is also worth mentioning that the general efflux pumps genes previously associated with an increased QAC tolerance (e.g., *mdrL* and *lde*) were detected in both QAC tolerant and sensitive isolates. The role of these genes may be explained by gene expression regulation differences between resistant and sensitive strains. It is possible that genetic variation in these genes cannot explain different QAC tolerance phenotypes, but rather that the differential expression regulation may be responsible for these effects.

Gene variants or specific genetic organizations located on plasmids, or a chromosome have been reported to have an effect on the disinfectant inhibitory concentrations. For instance, Dutta et al. (57) demonstrated that *L. monocytogenes* that carried *bcrABC* in different genetic locations, such as in categories VI (presumed chromosomal origin) and VII (composite transposon), exhibited lower MICs (30 mg/L) compared to categories I-V (MIC = 40 mg/L), which have been associated with plasmid origin. In the present study all *brcABC* carrying isolates showed the same tolerance to BC. Møretrø et al. (27) reported a variation among the tolerant phenotypes due to a non-synonymous mutation in the *qacH* gene. QacH harboring ^42^Ser exhibited higher MIC values than those carrying ^42^Cys. In this study, ^42^Ser was detected in QacH variants 4, 6 and 7, but no difference in the MIC values was observed. If further in-depth phenotypic differences could be investigated with a more sensitive method such as growth curve analysis (44).

Association between *qacH* location and its sequence has not previously been discussed in the literature, except for the report of a *qacH* homolog in a *L. monocytogenes* isolate from FPE in a German production facility, which lacked the other Tn*6188* genes (NG_076646.1 of isolate 16-LI00532-0) (22) and a recent report in a clinical isolate from Norway (35). In our study, we detected a *qacH* homolog in RepA G4 plasmid, with 100% nucleotide similarity to NG_076646.1 and 91-92% nucleotide identity to the chromosomal *qacH* (HG329628.1). The plasmid-borne *qacH* homolog was associated with a *tetR* gene, with 72% nucleotide identity to the chromosomal *tetR* located immediately upstream of *qacH* . Similarly, Chmielowska et al. (45) and Schmitz-Esser et al. (12) observed that the RepA G4 plasmids, although rarely found in their studies, encoded a putative novel QAC transporter. However, when we used the broth microdilution method to test the isolates, differences in MIC values to BC there were not difference in the tolerance of isolates with plasmid-borne *qacH* and the *qacH* on Tn*6188*. Additionally, it was seen that all RepA G4 plasmids had a complete transfer module, which suggests that they can be mobilized by a conjugative transfer. However, despite potential mobility, the occurrence of the RepA G4 plasmids tends to be low. In our study only seven isolates carried these plasmids, with similar low numbers observed by Chmielowska et al. (45) (n = 4), Schmitz-Esser et al. (12) (n = 8), one isolate in a study by Fagerlund et al. (35) (n = 1). Notably, five of the strains carrying the novel *qacH* homolog gene in the present study are from FPEs in three European countries (Italy, Spain and UK), isolated between 2015 and 2020.

Regarding the *bcrABC* cassette structures identified in this study, the *bcrABC* genetic environment consisted of genes encoding recombinase, glyoxalase and oxidoreductase, with exception of CC321 isolates which carried *bcrABC* on a composite transposon flanked by two IS*1216* elements. It is important to note that structure 8 in this study (Fig. S5) corresponds to category 7 in Dutta et al. (46). Contrary to Dutta et al., who observed both plasmid- and chromosome-located *bcrABC* cassettes, all structures in this study carried either the *cadC1* gene conferring resistance to cadmium, the mercury resistance operon, the stress response module NiCo riboswitch-*gbuC*-*npr* and/or the *clpL* gene, all suggesting plasmid origin of the contigs. None of the *bcrABC* genes in this study were associated with chromosomal origin. As all *bcrABC* structures were found to co-occur with other genetic elements conferring stress resistance, their impact on the persistence potential of *L. monocytogenes* in FPEs should be further investigated. Castro et al. (32) found that mobile genetic elements harbouring resistance genes for arsenic and cadmium were significantly more prevalent among persistent *L. monocytogenes* isolates from farm environments than in non-persistent isolates. Conversely, persistent isolates have been found to carry mutations associated with attenuated virulence, including a truncated *inlA* gene (47), which is in line with our findings of 98% of the FPE CC121 isolates possessing a PMSC in the *inlA* gene. Additionally, co-occurrence of QAC tolerance genes with other stress resistance genes, such as metal and antibiotic resistance genes, can contribute to their adaptation and persistence in FPEs. If exerted on disinfectant selective pressure, a response together with tolerance to other stressors may occur. It is common for BC tolerant isolates from food and food processing facilities to be cadmium resistant (48). However, our results also showed that in addition to cadmium resistance genes, the heat resistance gene *clpL*, plasmid replicon genes and the SSI-2 genes are strongly associated with QAC tolerant phenotype (Fig. 5A, Fig 6B). The SSI-2, involved in alkaline and oxidative stress responses, has been described to predominate in the hypovirulent CC121 isolates, mostly associated with FPE, and also carrying the *qacH* gene (49).

While not highly prevalent in our isolate collection, the high global prevalence of the *bcrABC* cassette and its predominance in phylogenetically distant CCs, may be due to its location on plasmids with various structures that allow for transfer between many different CCs (50). Evolutionary, the high prevalence could be due to an earlier acquisition of the cassette in the *Listeria* genome compared to the other QAC *L. monocytogenes* genes. Another possible explanation for the *bcrABC* predominance in food production facilities is their co-occurrence with other stress resistance genes, which can aid the survival of the isolates that carry them. It is also possible that they have better genomic compatibility with CCs that are likely to get established as persisters in FPEs, such as CC121, CC9, or in hypovirulent genomes that carry PMSCs, or other signature genetic characteristics associated with FPE.

On the other hand, the global differences in the gene prevalence as clearly seen for *emrE* (Fig. S7, S8) with much higher prevalence rates in some countries could be related to trading patterns between different countries/continents (e.g., US and Oceania/Australia) as contamination of raw materials with specific strains in one country can affect the *L. monocytogenes* prevalence rates in another country, as well as practices for application and use of disinfectants. The higher sequencing rates and public availability of bacterial genomes in US could partially explain the increased finding of *brcABC* among the global dataset in this study (Fig. S8).

The majority of the QAC tolerance genes found in the global dataset were associated with food production environment and food and feed samples, as previously reported by other studies (50,51), with the exception of *emrC* whose predominance in the production environment was merely 1.6%, compared to 44.2% seen in clinical samples. The latter is in agreement with the studies of Daeschel et al. (51) and He et al. (50), who examined the prevalence of QAC tolerance genes among *L. monocytogenes* isolates from US processing facilities and produce processing environments, respectively, and did not detect *emrC*.

Despite being associated with one of the most prevalent CCs within the FPEs, CC121 and CC9, the negligible detection of the *qacH* gene compared to the *bcrABC* cassette in the North American *L. monocytogenes* isolates, could possibly be due to a lower CC121 distribution in this geographic region or differences in the use of disinfectants in different countries/continents. It appears that the QAC-tolerant *L. monocytogenes*, which persist in the FPE in US, are non-CC121 *bcrABC*-harbouring isolates, such as CC5, CC321, CC155, CC7, CC9 and CC199 (35,41,46). Other large-scale studies comprising of *L. monocytogenes* isolates from Europe, report *qacH* as the predominant QAC tolerance gene, 18.8% in a French study (52) and 18.9% in an EU-study (53). This also leads to the idea that the distribution of QAC tolerance genes in different geographic regions is related to the predominance of CCs (51).

Despite its widespread distribution among geographic regions and CCs, *bcrABC* has never been associated with CC121, probably due to the high prevalence of *qacH* in this CC or possible incompatibility of two QAC tolerance genes in one *L. monocytogenes* genome. We also found 17 isolates that carried two different QAC tolerance genes, specifically *bcrABC* and *qacH*, *bcrABC* and *emrC*, and *qacH* and *emrC*. Kropac et al. (54) showed that *emrC* could be transformed in a *L. monocytogenes* CC2 genome carrying chromosomal *qacH*. The authors even reported slight increase in the BC MIC value of the transformant compared to the MIC value of the recipient cell and hypothesized that acquiring additional BC efflux pump gene can have an additive effect on the tolerance level. It is unknown, however, if in the nature it is advantageous for *L. monocytogenes* to harbour two QAC tolerance genes, especially the *emrC* gene which is located on a high copy plasmid. Similarly, Daeschel et al. (51) reported that 50% of the isolates screened in their study contained one or more of *bcrABC*, *emrE* or *qacH* genes, however, harbourage of more than one QAC tolerance gene in a single *L. monocytogenes* genome similarly to our study could be explained by their dataset, consisting of sequencing data downloaded from NCBI Sequence Read Archive that could contain contaminating reads.

In conclusion, this study showed that QAC tolerance can be *in-silico* predicted as genotype-phenotype concordance was 99% for QAC-based disinfectants for the four assessed QAC tolerance genes which all conferred the same level of tolerance to QACs as seen in the MICs. The propagation of QAC tolerance genes is however not evenly distributed globally with differences between continents and countries which showcases that understanding the development of *L. monocytogenes* disinfectant sensitivity and tolerance can be monitored using publicly available WGS data as different QAC gene variants and genetic contexts exist.

## MATERIALS AND METHODS

### Listeria monocytogenes isolates

A total of 1671 *L. monocytogenes* isolates recovered from human (n = 83; 5%), animal (n = 122; 7%), food (n = 839; 50%), FPE (n = 488; 29%), feed (n = 4; <1%), natural environment (n = 32; 2%), farm environment (n = 66; 4%) and unknown sources (n = 22; 1%) were included in this study. Additionally, 14 *L. monocytogenes* isolates (<1%) with unspecified clinical origin and one *L. monocytogenes* isolate (<1%) with unspecified environmental origin were also included. The isolates originated from 19 countries and were collected within a time span of 98 years, from 1924 to 2021 (Fig. 1C, Table S1).

### Determination of the *Listeria monocytogenes* minimum inhibitory concentrations to QACs

The minimum inhibitory concentrations (MIC) of *L. monocytogenes* to two pure biocide substances, benzalkonium chloride (BC) (500 g/L, Thermo Fisher, Kandel, Germany), didecyldimethylammonium chloride (DDAC) (500 g/L, Sigma-Aldrich, Denmark), and a commercial disinfectant - Mida SAN 360 OM (10-15% 2-methoxymethylethoxy propanol, 3-5% didecyl dimethyl ammonium chloride, < 3% 1,2-ethanediol; Christeyns, Denmark) were tested by an in-house optimized broth microdilution assay according to Wiegand et al. (55). The *L. monocytogenes* isolates were streaked from -80°C on trypticase soy agar plates (TSA; 40 g/L) and incubated at 37°C overnight. Three single colonies per isolate were transferred to 0.1x trypticase soy broth (TSB; 3 g/L) and cultured at 15°C for 48 h under stationary conditions until the optical density at 620 nm (OD620) measured ∼0.1, corresponding to 10^8^ CFU/mL. The final inoculum concentration was 10^5^ CFU/mL per well. The range of the disinfectant concentrations was selected according to previously reported MIC values (17) using two-fold dilutions. A positive control consisting of inoculated 0.1x TSB and a negative control consisting of sterile 0.1x TSB were included in each 96-well plate. The plates were sealed with adhesive film (ThermoFisher, Denmark) and incubated at 15°C for 48 h. The MIC, defined as the lowest biocide concentration at which the *L. monocytogenes* growth was inhibited, was determined by measuring the *L. monocytogenes* OD620 by a Multiscan FC Microplate Reader (Thermo Scientific, Denmark). The threshold for growth was set at OD620 ≥ 0.08, which is 60-65% of OD620 of the positive control and 200% of the negative control. The experiment was performed in two independent biological replicates with three technical replicates each. The result was considered valid if two out of the three technical replicates had identical MIC values. One two-fold MIC variation between biological replicates was considered acceptable and the higher MIC value was reported as the final result, unless the two-fold dilution difference was at the cut-off for tolerance (MIC ≥ 1.25 mg/L). In this case, the test was repeated a third time.

### DNA extraction and whole-genome sequencing

For whole-genome sequencing (WGS), as part of the current study, the *L. monocytogenes* isolates (n = 1,244) were grown on TSA at 37°C, overnight. A single colony per strain was transferred to 1.8 ml TSB and grown at 37°C overnight. Genomic DNA was extracted by the DNeasy Blood and Tissue Kit (Qiagen, Denmark) following the manufacturer’s recommendations except that the DNA was eluted in 10 mM Tris-HCl (pH = 8.5) (BioNordika, Denmark). The DNA concentration was measured using Quant-iT dsDNA high sensitivity kit (Invitrogen, Denmark) by VICTOR X2 Multilabel Microplate Reader (Spectralab Scientific Inc.). Sequencing libraries were constructed using the Nextera XT Library Prep Kit (Illumina, San Diego, CA, USA), normalized and denatured for loading in a NextSeq 500/550 Mid Output v2.5 Kit (300 cycles) (Illumina), and pair-end sequenced on a NextSeq 500 platform (Illumina). The remaining 427 *L. monocytogenes* isolates were previously sequenced (Table S1).

### Assembly, species identification and *in silico* sub-typing

The raw sequencing data of the 1,671 isolates were processed with the FoodQCpipeline v1.6 (https://bitbucket.org/genomicepidemiology/foodqcpipeline/src/master/), which uses bbduk2 from bbtools (Bushnell B. sourceforge.net/projects/bbmap/) for trimming and SPAdes (56) for genome assembly. Quality control of the sequencing reads was performed before and after trimming by FastQC v0.11.5 (57). Quality of the assemblies was assessed by Quast v4.5 (58) and thresholds for number of contigs (≤ 300 contigs) and genome size (3 Mb ± 0.5 Mb) were established. *In silico* species identification was performed using KmerFinder v2.0 (59). Assemblies were submitted to the BIGSdb-*L. monocytogenes* Pasteur MLST database (https://bigsdb.pasteur.fr/listeria/) for sub-typing (Table S1).

### Core-genome MLST and phylogenetic analysis

The assemblies were annotated with Prokka v1.14.6 referencing the genus *Listeria* and the species *monocytogenes* (60). Core-genome alignment was generated by Roary v3.13.0 using the .gff files from Prokka as input, with 95% blastp identity threshold and paralog splitting (-s) disabled (61) to prevent presumed paralogous genes from being split into different gene groups. The core-gene alignment was trimmed with trimAI (62) and option -gappyout to decide optimal thresholds based on the gap percentage count over the whole alignment. A maximum-likelihood (ML) phylogenetic tree was inferred using IQ-TREE v2.1.3 with the GTR+G nucleotide substitution model and 1000 bootstrap replicates (--ufboot 1000) and mid-point rooted. The trees were visualized and annotated in iTOL (63) or ggtreeExtra R package (64).

### Identification of plasmids

The plasmid replicon gene screening was carried out by BLAST+ v2.2.31 (65) with a database consisting of representative *repA* genes from each of the eleven phylogenetic groups identified by Chmielowska et al. (2021) (66), the *repA* G12 by Fagerlund et al. (2022) (35), the *repB* genes from the small *Listeria* spp. plasmids, and the complete sequences of the plasmids with an unknown replication system (Table S3). The blastn was performed with the fasta files as queries against the *repA* database with minimum identification (--minid) and coverage (--mincov) thresholds of 80 and 95, respectively, in ABRicate v1.0.1 (Seemann T, Abricate, Github https://github.com/tseemann/abricate). The *repA*- harboring contigs were extracted from the assemblies using *awk* and the *repA* gene sequences subsequently extracted by getfasta in bedtools v2.30.0 (67). Translated *repA* gene sequences were aligned with MAFFT v1.5.0 and used to infer maximum likelihood tree in IQ-TREE using the Le and Gascuel (LG) amino-acid substitution model and ultrafast bootstrapping (--ufboot 1000). Reference RepA sequences from each of the twelve RepA groups (G1-G12) (35,45) were included in the tree. The tree was visualized and annotated in iTOL.

### Genetic organization of the *bcrABC*-harbouring plasmids

Plasmid contigs carrying *bcrABC* were annotated by Prokka with --compliant option. The contigs were aligned in Geneious Prime 2023.0.4 by MAUVE and MAFFT and grouped according to their genetic organization. From each group of identical contigs (indels or SNPs were ignored), one contig was chosen as a representative and the .gbf from Prokka uploaded to Clinker (68) for pairwise alignment and visualization. Plasmid contigs from each *bcrABC* category were aligned to a custom database consisting of complete *Listeria* spp. plasmids carrying the *bcrABC* cassette (45,69). Additionally, the plasmid contigs from each category were compared with publicly available sequences in NCBI.

### Virulence and resistance genes profiles

A set of 97 virulence and resistance genes identified based on the literature (Table S4) were searched in the *L. monocytogenes* genomes using a custom database in ABRicate with minimum identification (--minid) threshold of 90%. Genes detected on contigs with very low coverage (<2×) compared to the rest of the contigs in each assembly were excluded as possible contamination. Except for truncations/interruptions in *actA*, *inlA*, *inlB* and *comK* genes, a gene was considered present when coverage was >90%. Additionally, the assemblies were screened against the ResFinder v4.0 database for the presence of antimicrobial resistance (AMR) genes. Mutations and internal truncations in *inlA* and *inlB* genes were identified by extracting the genes as explained for plasmid replicon genes, translated into protein sequences and aligned to the EGD-e *inlA* (NC_003210.1:454534..456936) and *inlB* (NC_003210.1:457021..458913) as reference genes.

### *In silico* screening for disinfectant resistance genes in global *L. monocytogenes* isolates

To study the global prevalence of the genes associated with increased tolerance to QACs (*emrC*, *emrE*, *bcrABC* and *qacH*) in *L. monocytogenes*, publicly available raw sequencing data deposited in ENA as of 26 November 2018 were screened using COBS (COmpact Bit-sliced Signature index) v0.1.2 with default settings for *emrE* (NC_013766.2:c1850670-1850347), *emrC* (MT912503.1:2384-2770) *bcrABC* (JX023284.1) sequences and homology reduced to maximum 90% nucleotide identity for *qacH* variants (Table S1). COBS consisted of 661,405 assembled and indexed bacterial genomes and 26,006 of them were annotated as *L. monocytogenes* (71). The STs of the *L. monocytogenes* isolates positive for QAC tolerance genes were obtained from the metadata associated with COBS. Furthermore, the pair-end sequencing runs deposited in ENA between 27 November 2018 and 29 April 2021 were downloaded and screened for QAC tolerance genes using KMA with minimum template identity of 90% (72) . To index the database of biocide genes, a k-mer = 16 was used as a default. The STs of the *L. monocytogenes* isolates harboring the disinfectant resistance genes were determined by stringmlst v0.6.3 (73) with k-mer = 35 using *L. monocytogenes* MLST database and converted to CCs using the Pasteur’s BIGSdb-Lm MLST database (https://bigsdb.pasteur.fr/). The world map for QAC tolerance gene distribution was produced using the R packages *mapPies* (https://search.r-project.org/CRAN/refmans/rworldmap/html/mapPies.html) and *rworldmap* version 1.3-6 (https://cran.r-project.org/web/packages/rworldmap/rworldmap.pdf).

### Statistical analysis

The heterogeneity in proportion of clonal complexes (CCs), geographical locations and isolation sources that were positive within and between genes was estimated using a random-effects model proposed by and implemented in the R package meta to produce forest plots (74). Statistical heterogeneity within and between groups was estimated using the Cochran chi-square test and the Cochrane I^2^ index. The Pearson’s chi-squared association test was performed using Microsoft Excel to determine statistically significant (p < 0.05) association between presence or absence of plasmids, stress survival and virulence genes in the phenotypes defined as tolerant or sensitive to BC.

### Data availability

The raw sequencing data have been deposited in the European Nucleotide Archive with metadata overview in Table S1.

## ACKNOWLEDGEMENTS

The authors gratefully acknowledge Pia Engelsmann, Resadije Idrizi, Rannvá Høgnadóttir Houmann (Research Group for Food Microbiology and Hygiene, DTU) and Jacob Dyring Jensen, Gunhild Larsen and Christina Aaby Svendsen (Research Group for Genomic Epidemiology, DTU) for excellent technical assistance throughout the project. Stephanie Brown (Food Innovation Center, Oregon State University, Portland, OR, USA) is thanked for her assistance with some of the US *L. monocytogenes* isolates. Lone Gram (Department of Biotechnology and Biomedicine, DTU) is kindly acknowledged for sharing *L. monocytogenes* isolates for this project. The Institut Pasteur teams are acknowledged for the curation and maintenance of BIGSdb-Pasteur databases at https://bigsdb.pasteur.fr/. This work was supported by the Danish Dairy Research Foundation, the Milk Levy Fund and Karl Pedersen og Hustrus Industrifond (DI-2019-07020) grants.

## CONFLICTS OF INTEREST

The authors have no conflicts of interest to declare.

## SUPPLEMENTARY FIGURES

**Figure S1.**
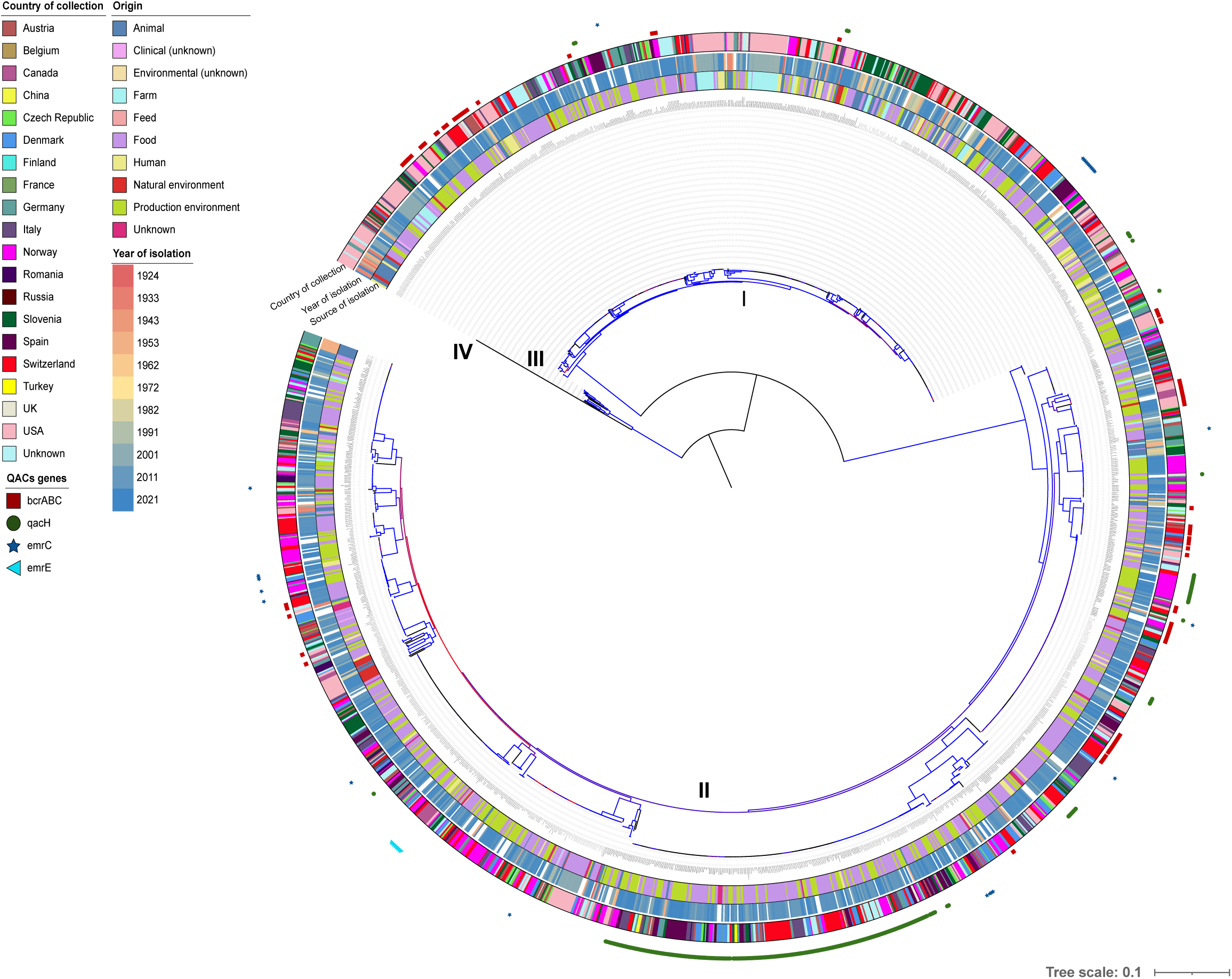
Mid-rooted maximum likelihood (ML) tree constructed from 2.19 Mb core-genome alignment using IQ-TREE with 1000 ultrafast bootstraps and GTR+G nucleotide substitution model. The color of the branches represents bootstrap support values, from 50 (red) to 100 (blue). Roman numbers represent the *L. monocytogenes* phylogenetic lineages. The rings from the inner to outer direction represent the source of isolation (origin), year of isolation (if known) and country of collection. Presence of QAC tolerance genes is presented by symbols outside the rings.

**Figure S2.**
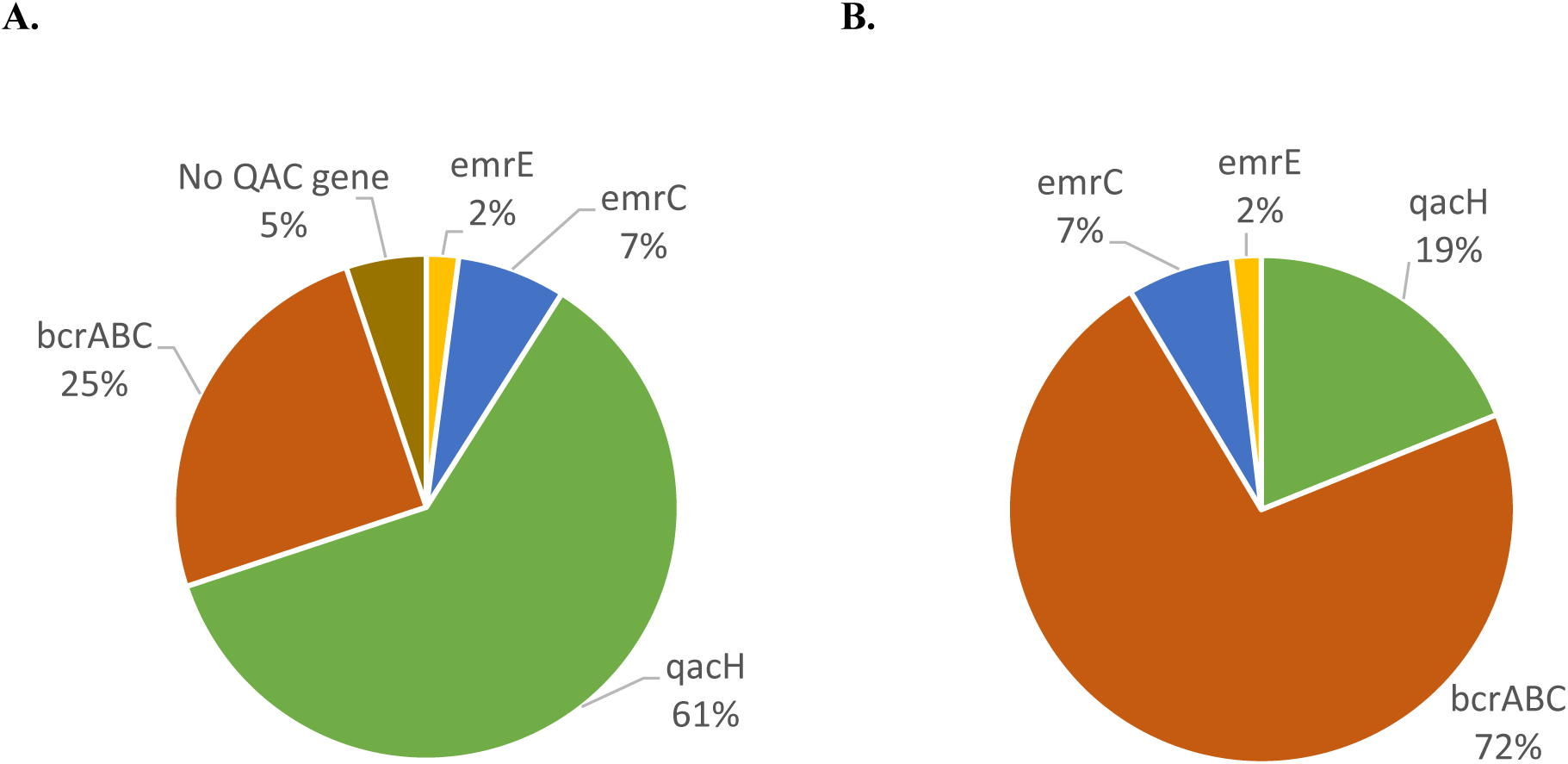
Distribution of the QAC tolerance genes in the 388 QAC-tolerant *L. monocytogenes* isolates in this study (A) and in the ENA *L. monocytogenes* isolates as of April 2021 (B).

**Figure S3.**
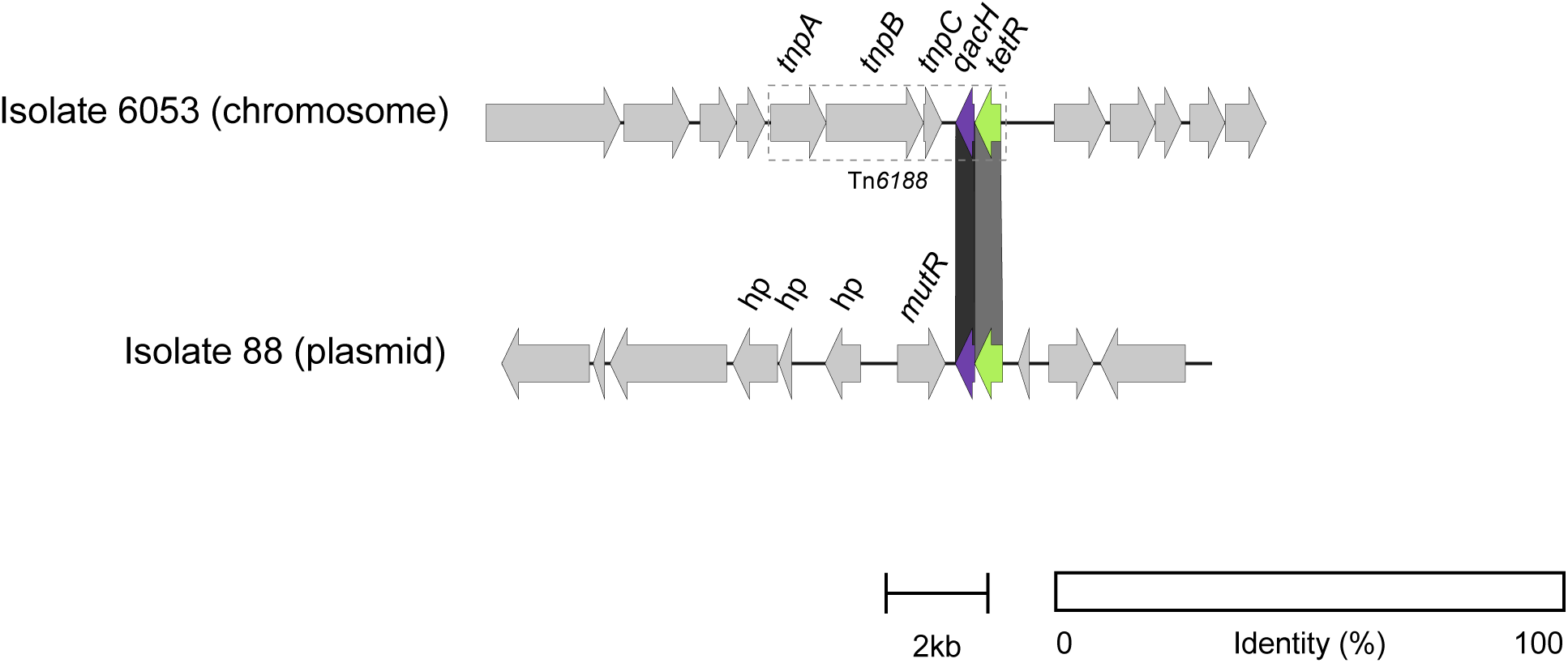
Genetic organisation of the *qacH* gene located on the chromosome and plasmids. The chromosome-located *qacH* is part of the previously described Tn*6188* (19), while plasmid-located *qacH* has different genetic environment, consisting of *tetR* (72% nucleotide identity to *tetR* on the chromosome), *mutR* transcriptional regulator and genes encoding hypothetical proteins. The nucleotide identity between plasmid- and chromosome-located *qacH* was 91%.

**Figure S4.**
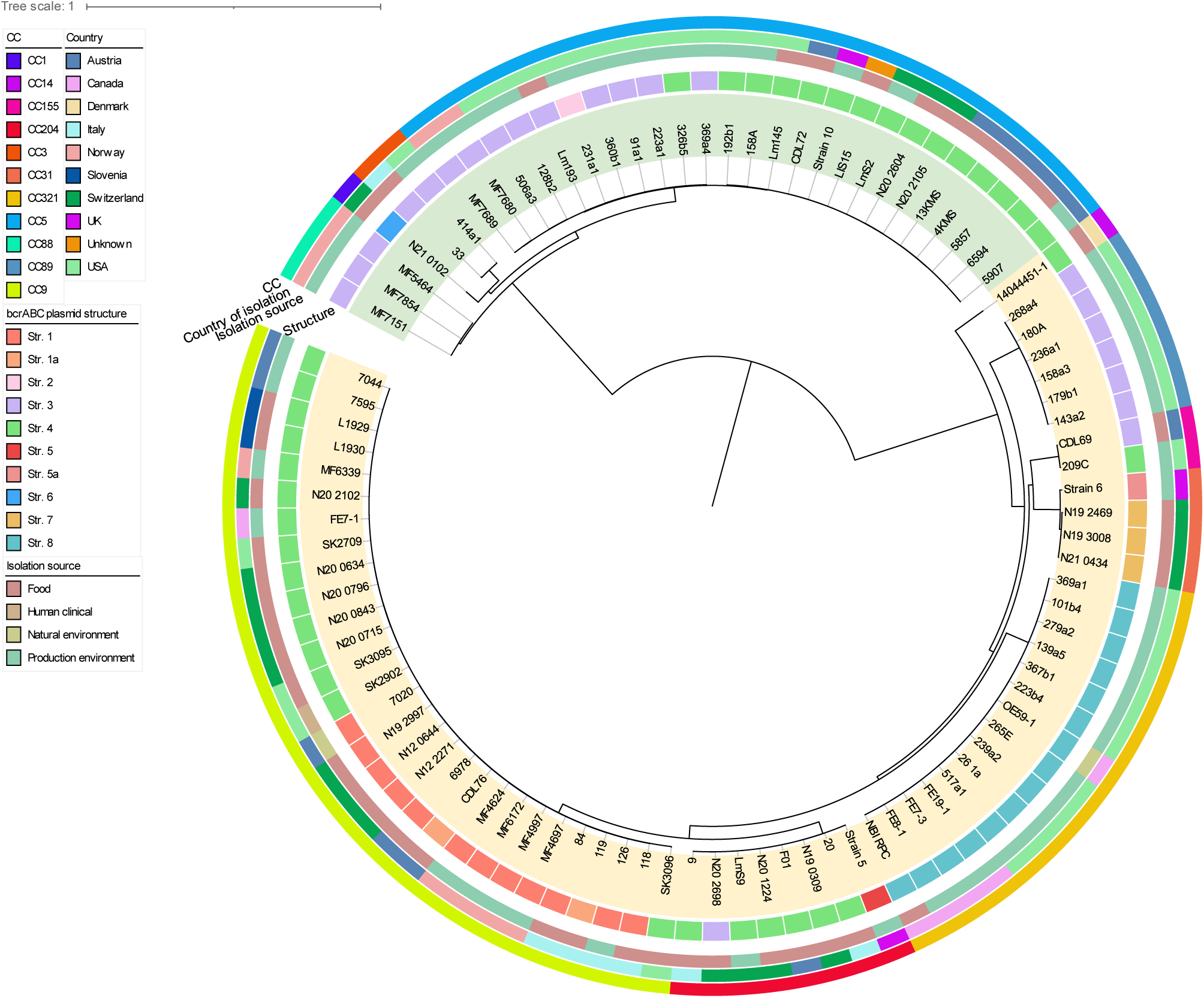
Mid-point rooted SNP-based phylogenetic tree of the 96 *bcrABC*-containing *L. monocytogenes* isolates.

**Figure S5.**
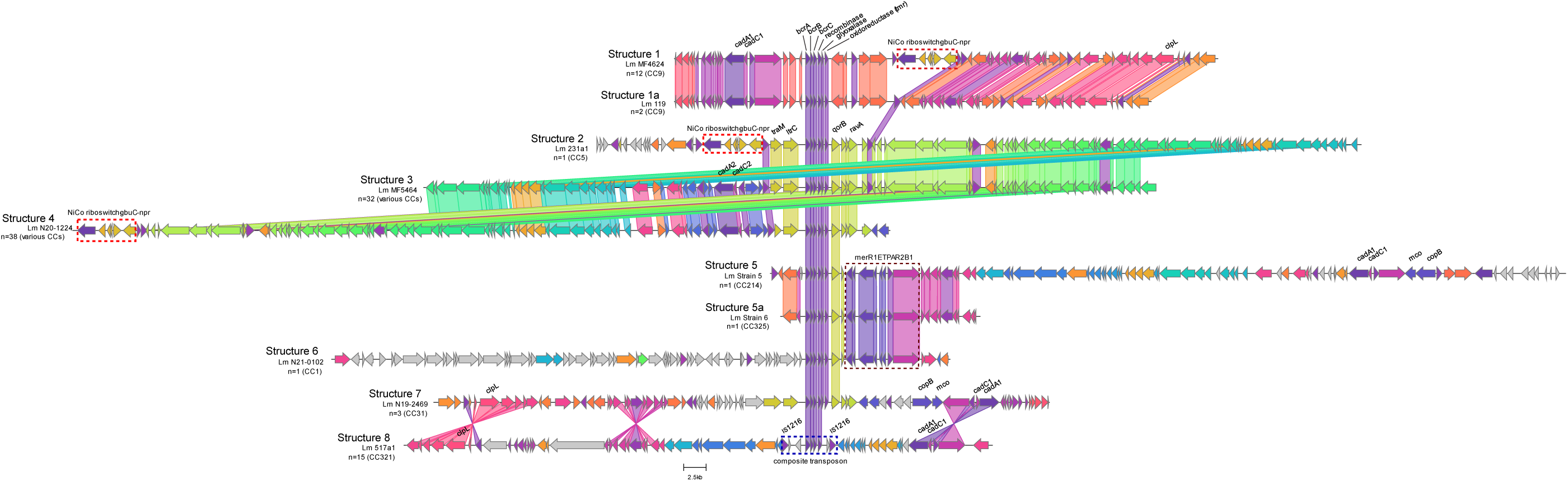
Organisation of the *bcrABC*-harbouring contigs grouped into structures according to their genetic contexts. Links between homologous genes are shown in different colours (the threshold for percentage identity is 90%). Genes in grey have no homologous genes within the analysed plasmid contigs. The red dashed rectangles represent a mobile genetic element carrying Nico riboswitch, *gbuC* and *npr* genes, and the blue dashed rectangle represents the composite transposon in plasmid structure 8. All stress resistance genes identified by blastn are annotated in the contigs; “n” indicates the number of isolates in each structure. The genes are automatically coloured by clinker according to their protein functions

**Figure S6.**
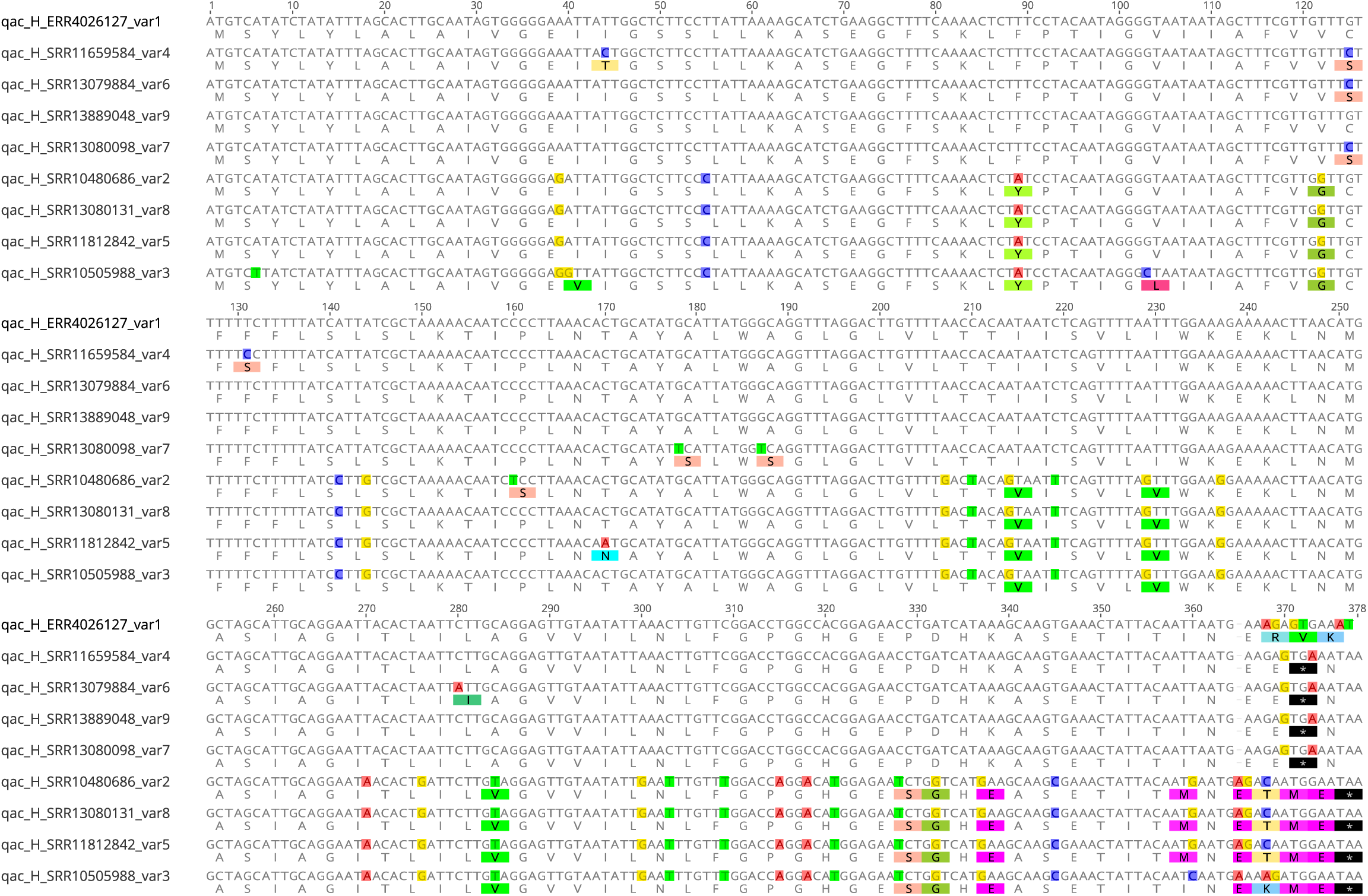
Nucleotide alignment and translation of the nine *qacH* variants identified among the global *L. monocytogenes* dataset with nucleotide identity above 90%. The ENA run accession numbers are given for each isolate harbouring the respective variant.

**Figure S7.**
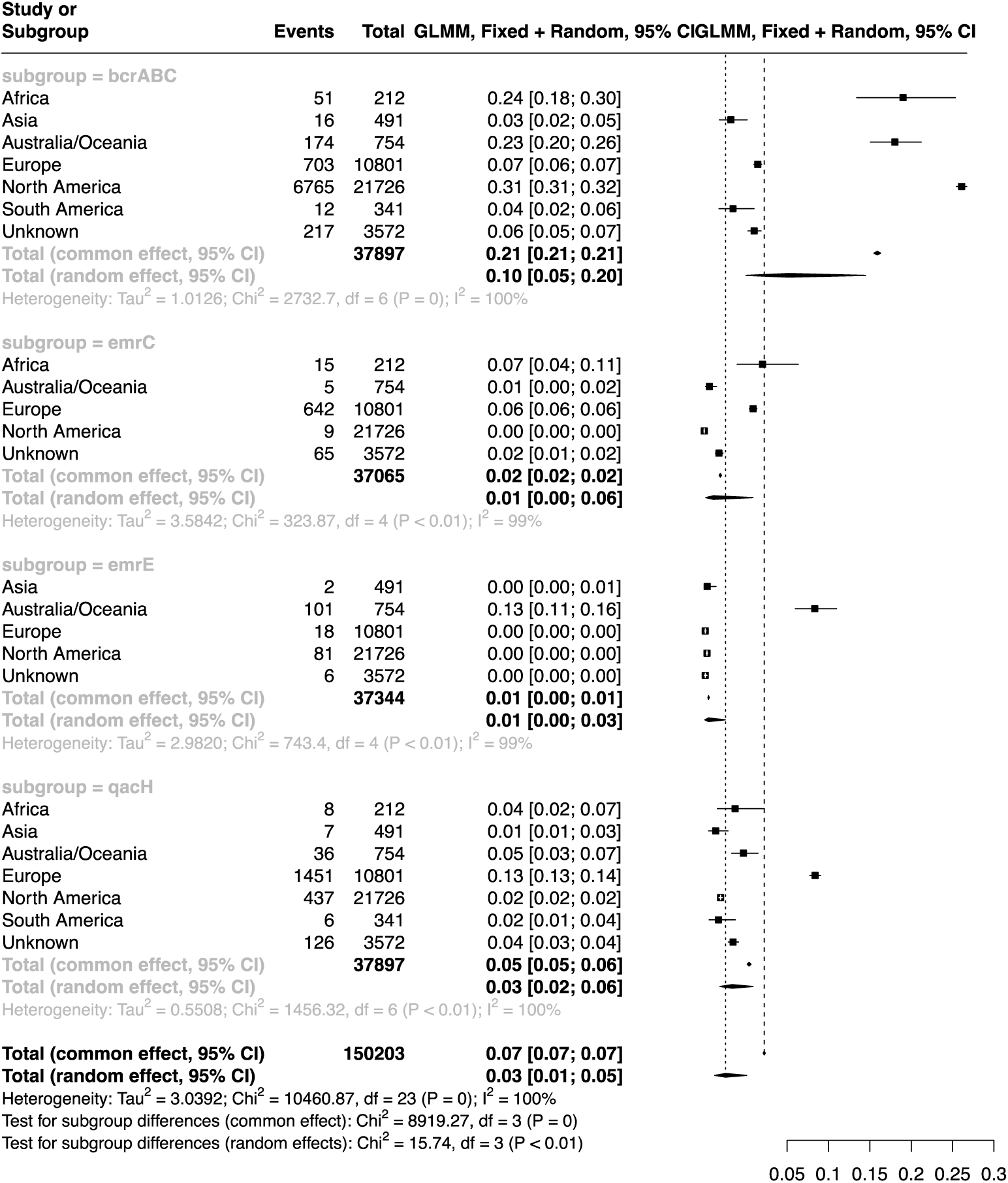
Forest plot showing the association between QAC tolerance genes and continent in the global *L. monocytogenes* dataset.

**Figure S8.**
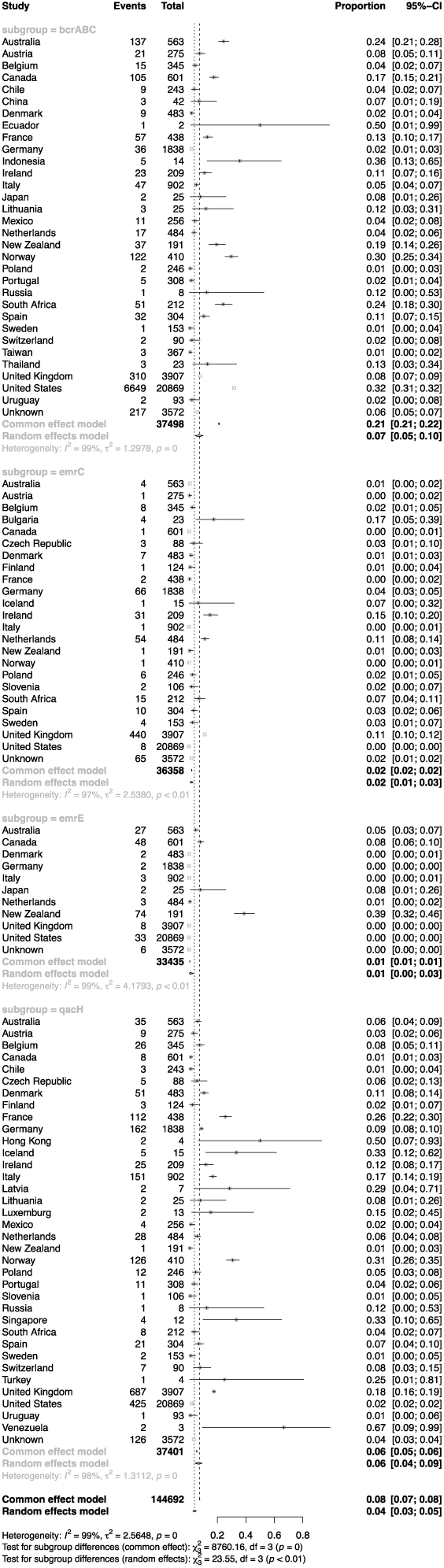
Forest plot showing the association between QAC tolerance genes and country of isolation in the global *L. monocytogenes* dataset.

**Figure S9.**
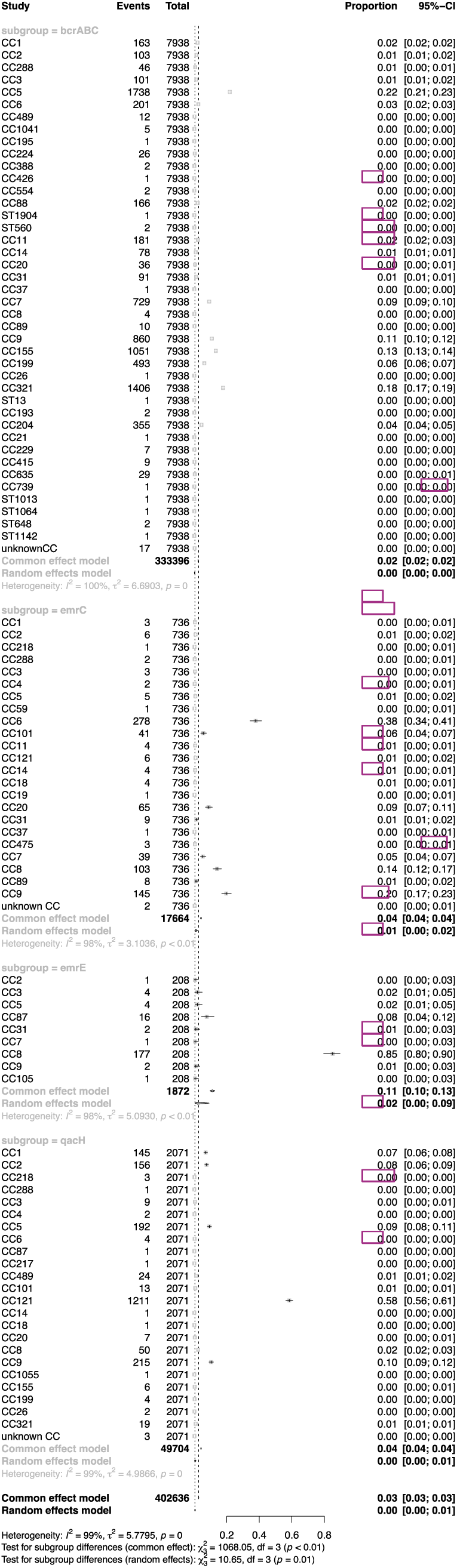
Forest plot showing the association between QAC tolerance genes and CC in the global *L. monocytogenes* dataset.

**Figure S10.**
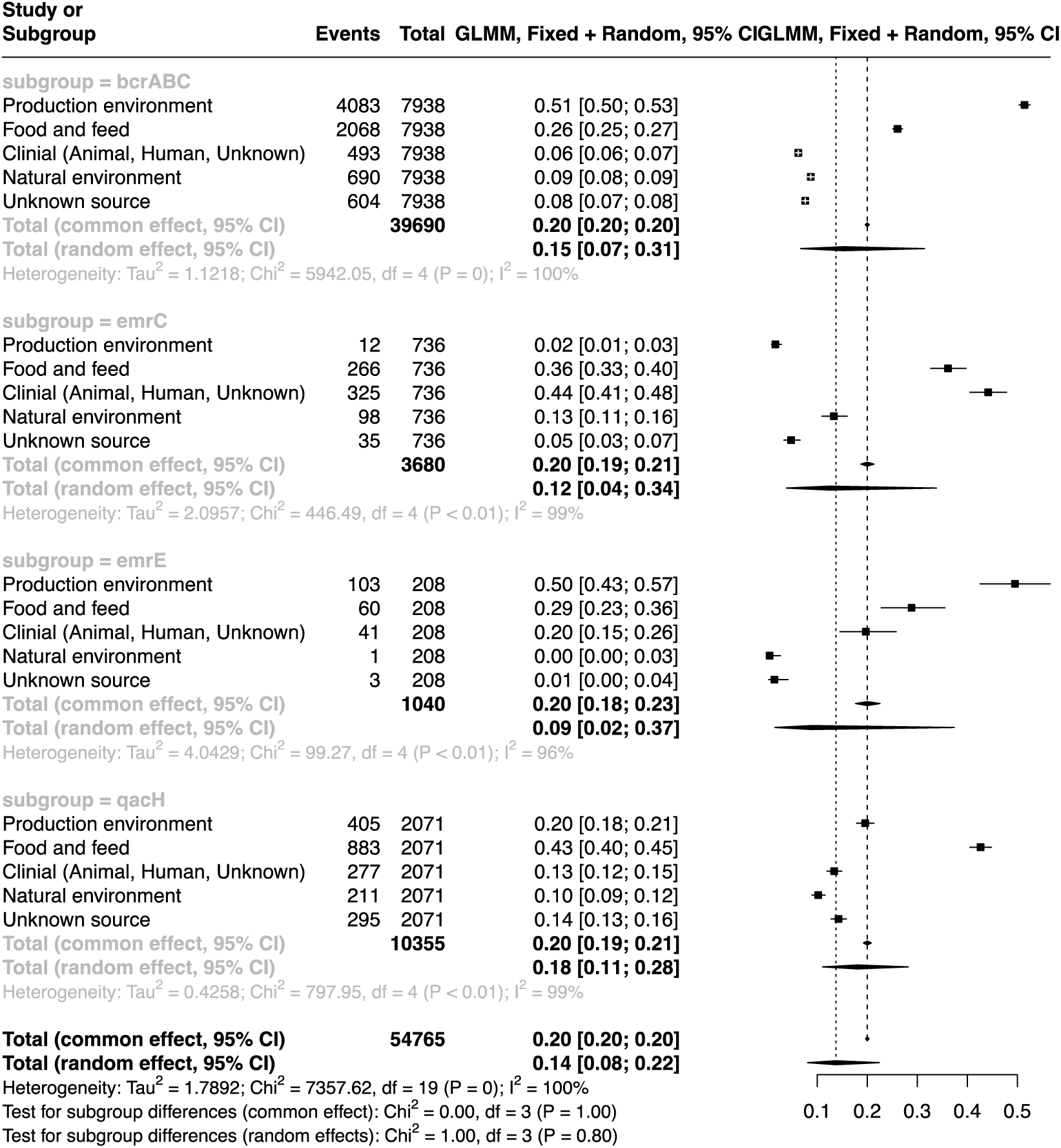
Forest plot showing the association between QAC tolerance genes and source of isolation in the global *L. monocytogenes* dataset.

